# Proteomic Signatures of Microbial Adaptation to the Highest UV-Irradiation on Earth: Lessons from a Soil Actinobacterium

**DOI:** 10.1101/2021.07.11.451938

**Authors:** Federico Zannier, Luciano Raúl Portero, Thierry Douki, Wolfgang Gärtner, María Eugenia Farías, Virginia Helena Albarracin

## Abstract

In the Puna region, the total solar irradiation and the UV incidence is the highest on Earth, thus, restraining the physiology of individual microorganisms and the composition of microbial communities. UV-resistance of microbial strains thriving in High-Altitude Andean Lakes was demonstrated and their mechanisms were partially characterized by genomic analysis, biochemical and physiological assays. In this work, we present the molecular events involved in the adaptive response of the model HAAL extremophilic actinobacterium *Nesterenkonia* sp. Act20 under artificial UV-B radiation, herein called as UV-resistome. Proteomic profiles of cultures exposed to different UV-experimental conditions showed that the leading systems for adaptation to the UV-challenge *in-vitro* are DNA repair and antioxidant mechanisms.

## INTRODUCTION

Free-living microorganisms endure physicochemical stochasticity of their environments, and due to their size and simplicity, they have relatively limited capabilities to adapt to or avoid the exposure of a high level of harmful environmental agents. Therefore, in unicellular organisms, genetic regulation plays a pivotal role to survive under stress conditions as it contributes to rescheduling gene expression and coordination of the synthesis of defense proteins, usually on the expense of expression of genes related to the cellular growth system. Changes in the concentration and activity of proteins encoded by these genes constitute the ‘adaptive response’ that, in turn, establishes a feedback circuit that modulates its intensity and duration, allowing the genetic system to restore the levels of expression before the onset of the stimulus or to adapt to the new environmental condition (Aertsen & Michiels, 2004; Kærn et al., 2005; Ramírez Santos et al., 2001). Thus, the adaptive response is a vital element for microorganisms that live in habitats with large physicochemical fluctuations allowing them to anticipate and acclimatize to harmful conditions (Thattai & Van Oudenaarden, 2004).

In the Central Andean region in South America, high-altitude ecosystems (3500-6000 m asl) are distributed across Argentina, Chile, Bolivia and Peru, in which poly-extremophilic microbes thrive under extreme environmental conditions (Albarracín et al., 2009; Albarracín, Gärtner, et al., 2016; Farías, 2020; Orellana et al., 2018; Vignale et al., 2021). Previous works proved the high tolerance profiles of these extremophiles to antibiotics, dryness, heavy-metal, arsenic, hypersalinity, and UV irradiation (Albarracín et al., 2009, 2012; Alonso-Reyes, Farias, et al., 2020; Belfiore et al., 2013; Dib et al., 2009; Flores et al., 2009; Kurth et al., 2015, 2017; Omar Federico Ordoñez et al., 2018; Orellana et al., 2018; Perez et al., 2020; Pérez et al., 2017a; Portero et al., 2019; Nicolas Rascovan et al., 2015; Rasuk et al., 2017; Saona et al., 2021; Zannier et al., 2019; Zenoff, Siñeriz, et al., 2006). Moreover, various underlying mechanisms and cellular processes were identified that indicate how these indigenous microbes cope with these multiple stress conditions, thus presenting an interesting side-aspect of opening avenues of research for novel biotechnological applications.

In the Puna region, total solar irradiation and the UV incidence is the highest on Earth, setting strong limits to the physiology of individual microorganisms and the composition of microbial communities in this region (Albarracin et al., 2011; Cabrol et al., 2004, 2009; Escudero et al., 2007; Flores et al., 2009; Omar F. Ordoñez et al., 2009; Pérez et al., 2017b; Nicolás Rascovan et al., 2016; Toneatti et al., 2017; Zenoff, Siñeriz, et al., 2006; Zenoff, Heredia, et al., 2006). There is extensive research on these poly-extremophiles focusing on the influence of UV irradiation on the molecular profiles and adaptive strategies of model microorganisms isolated from shallow lakes and soil across different sites in the Argentinean and Chilean Central Andes (Albarracin et al., 2011, 2015; Albarracín, Gärtner, et al., 2016). Genomics and ultrastructural and physiological assays identified a number of UV-resistance mechanisms of bacterial and archaeal strains (Albarracín et al., 2009, 2012; Albarracín, Kraiselburd, et al., 2016; Alonso-Reyes, Farias, et al., 2020; Alonso-Reyes, Galván, et al., 2020; Flores et al., 2009; Kurth et al., 2015; Portero et al., 2019; Rasuk et al., 2017; Toneatti et al., 2017; Zenoff, Heredia, et al., 2006; Zenoff, Siñeriz, et al., 2006).

Andean microbes’ high UV-resistance profile points to the existence of a network of physiological and molecular mechanisms triggered by ultraviolet light exposure. We called this “UV-resistome” (Kurth et al., 2015; Portero et al., 2019, Alonso-Reyes et al., 2020) that includes some or all of the following subsystems: (i) UV sensing and effective response regulators, (ii) UV-avoidance and shielding strategies, (iii) damage tolerance and oxidative stress response, (iv) energy management and metabolic resetting, and (v) DNA damage repair. Genes involved in the described UV-resistome were recently described in the genome of *Nesterenkonia* sp. Act20, an actinobacterium which showed survival to high UV-B doses as well as efficient photorepairing capability (Portero et al., 2019).

In this work, a proteomic approach dissects the adaptive response of *Nesterenkonia* sp. Act20 against UV-B radiation. The molecular events during the UV challenge and after photorepairing coincided with most hypothesized UV-resistome subsystems, especially the DNA repair tool kit. Our results demonstrate that UV-B exposure induces over-abundance of an exclusive set of proteins while recovery treatments tend to restore the proteomic profiles present before the UV exposure.

## MATERIALS AND METHODS

### Strains and growth conditions

*Nesterenkonia* sp. Act20 (Act20) belongs to the LIMLA extremophilic strain collection (Strain Number P156, PROIMI-CONICET). Axenic glycerol-freeze cultures were aerobically activated in a growth medium designed explicitly for this strain called “H medium” (i. e., <u>H</u>alophilic medium, 10 g NaCl, 3 g sodium citrate, pH 7-7.2) and cultured at 30 °C with agitation (220 rpm) overnight. *Nesterenkonia halotolerans* DSM 15474 was grown under the same conditions as Act20 and was used as a control strain to re-evaluate the artificial UV tolerance profile of Act20. Cultures of both strains were maintained in H agar (1.5%) for further inoculations.

### UV-lethal doses, acclimatization, and viability assays

Strains were activated in 20 ml of liquid culture medium and cultured overnight at 30 °C at 220 rpm. The cells were then centrifuged at 7000 rpm for 5 minutes, and the pellets were suspended in physiological solution at OD_600_ corresponding to 10^6^-10^8^ CFU/ml (OD_600_ 0.60 for both *Nesterenkonia* sp. strains). Serial dilutions (1:10) of the cell suspensions were made for each strain, and 10 µl aliquots of each dilution were linearly plated onto the agar culture medium and left to air-dry for 15 minutes in the laminar flow chamber. Two plates were immediately incubated in the dark (DR Control) or under artificial white light (FR Control) conditions, respectively, making up the experimental controls for both the “Lethal Dose” (LD) (continuous exposure) and “Acclimatization” (Acc) (discontinuous exposure) assays. Then, for the lethal dose test, eight plates were exposed in pairs for 15, 30, 60, and 90 minutes to artificial UV-B irradiation at 5.4 W/m² (4.86 kJ/m², 9.72 kJ/m², 19.44 kJ/m² and 29.16 kJ/m²) using a Vilbert Lourmat VL-4 lamp (maximum intensity at 312 nm). Irradiance was quantified with a radiometer (09811-56, Cole Parmer Instrument Company). After the end of each exposure period, one plate was incubated in darkness (DR), the other one was incubated under artificial photoactive light (FR, 72 hours for both strains). The acclimatization test consisted of exposing the samples to successive 15-minutes cycles of UV-B radiation at 5.4 W/m² alternating with 45-minutes of recovery intervals of incubation in the dark (DR 15’x45’) or under white light (FR 15’x45’), respectively. Thus, to achieve cumulative irradiation values comparable to the lethal dose test, the Acc plates were exposed to 1, 2, 4, and 6 cycles, respectively. Once each pair of Acc plates completed the assigned number of cycles, they were incubated under PAR light or dark for 72 hours, respectively. In both tests, plates were seeded in duplicate. At the end of the assay, plates were photographed and compared by eye.

### Preparation of cultures for photoproducts measurements and proteomic analysis

For this assay, *Nesterenkonia* sp. Act20 and *N. halotolerans* culture were exposed to the following experimental conditions; the treatment named “UV” consisted of cell suspensions in NaCl 0.9% (20 ml, OD_600_ ∼0.6), exposed to artificial UV-B irradiation (5.4 W/m² UV-B) for 20 min in quartz tubes. This dose was used to reduce 50% of the Act20 population. Treatments named “photorecovery (“FR”)” and “dark recovery (“DR”) were applied to cell suspensions exposed to UV as before, but then incubated under white light or in the dark for 120 min. In the treatment called “total dark control (“Dt”), cell suspensions were left in a quartz tube and kept in the dark during the UV-exposure and the recovery treatments. We use two other controls for the photoproducts measurements: non-exposed cell cultures at time 0 (T0) and non-exposed cell cultures after 20 min of incubation (T20).

All suspensions were then centrifuged, washed with NaCl (0.9%) and resuspended in 0.1 M Tris-HCl buffer, pH 7. The samples were stored at -70 °C until subsequent processing. The above-described protocol was carried out in triplicate, obtaining three independent biological replicates for each treatment.

### Photoproduct quantification

10 mL of cell suspensions of the different treatments T0, Dt, UV, FR, and DR conditions were centrifuged at 3000 g for 10 minutes at 4 °C. A cell suspension without exposure to any stimulus named T0 (initial time) was used as an additional control. Pellets were harvested and washed twice with distilled water. Total genomic DNA extraction was performed using a commercial genomic DNA kit (DNeasy Blood & Tissue Kit, Qiagen). Photoproducts were quantified using a pre-optimized procedure (Thierry Douki, 2013). After extraction, DNA was solubilized in an aqueous solution containing 0.1 mM desferrioxamine mesylate and then enzymatically hydrolyzed by incubation with nuclease P1, DNAase II, and phosphodiesterase II (2 h, 37 °C, pH 6), followed by the second stage of digestion involving phosphodiesterase I and alkaline phosphatase (2 h, 37 °C, pH 8). The digested DNA samples were injected into an Agilent 1100 Series HPLC system equipped with a reversed-phase ODB Uptisphere column (2 x 250 mm ID, particle size 5 μm, Interchim, Montluçon, France). The mobile phase (flow rate 0.2 ml/min) was an acetonitrile gradient (from 0 to 20 %) in a 2 mM aqueous solution of triethylammonium acetate. The HPLC flow was split and funnelled into an API 3000 electrospray triple quadrupole mass spectrometer operating in negative ionization mode. The pseudomolecular deprotonated ion of each photoproduct was collected and fragmented. Specific daughter ions of each photoproduct were quantified. Calibration curves were performed using proper reference compounds of varying concentrations. The results were expressed as the number of photoproducts per 10^6^ DNA bases. The standards for HPLC-ESI-MS / MS were synthesized according to a previously published procedure (T. Douki & Cadet, 2001). In summary, dinucleoside monophosphates were prepared by the triester synthesis. CPDs were obtained by photosensitized triplet energy transfer using acetophenone and UV-A treatment. The 6-4PPs were prepared by photolysis with UVC irradiation, and a subsequent UVA irradiation of these latter compounds produced the Dewar valence isomers. All photoproducts were HPLC purified. The four *cis-syn* cyclobutane pyrimidine dimer (CPDs) were measured, (thymine-thymine (TTCPD), thymine-cytosine (TCCPD), cytosine-thymine (CTCPD) and cytosine-cytosine sites (CCCPD)), with cytosine under its deaminated form uracil. Pyrimidine (6-4) pyrimidone photoproducts (6-4PP) at thymine-thymine (TT64) and at thymine-cytosine (TC64) sites, together with their Dewar valence isomers (DEWTT and DEWTC, respectively), were also quantified. The above-described protocol was carried out as four individual set-ups, obtaining four independent biological replicates for each treatment and each photoproduct category. Significant differences in photoproducts absolute concentrations and mean efficiency on photoproducts repair were analyzed through an ANOVA model and TukeyHSD test. Photoproducts data is available in supplementary file 3.

### Protein extraction protocol

Once the necessary three replicates of each treatment were obtained, we performed the protein extraction protocol described by (Dávila Costa et al., 2015). The pellets were washed with Tris buffer (25 mM), pH 7, EDTA (2 mM) and then resuspended with the same buffer composition, supplemented with 5 µl of a reducing solution (DTT 200 mM, Tris 100 mM, pH 7.8). The intracellular proteins were obtained by breaking the cells with a French press and incubating for 1 hour at room temperature to achieve complete denaturation. The protein concentration was measured using the Bradford assay. For every 50 µg of protein, 20 µl of reducing solution and 20 µl of alkylating solution (iodoacetamide, 200 mM, Tris, 100 mM, pH 7.8) were added to each sample. The mixture was allowed to incubate for 1 hour at room temperature and was then centrifuged at 16,100 g for 30 min (4 °C). Then, the proteins were precipitated with 10% TCA and incubated overnight (−20 °C). Samples were then centrifuged (16 100 g, 30 min, 4 °C), and the protein pellet was washed twice by rinsing with 500 ml pre-chilled (−20 °C) acetone. Once dried, the protein pellets were dissolved in ammonium bicarbonate (50 mM) and digested with trypsin at 37 °C for an incubation period of 14-16 hours. Finally, after digestion with trypsin, the concentration of peptides (10 µg/μl) was determined, and the samples were stored at -80 °C until further analysis by mass spectrometry (MS).

### Protein identification and mass spectrometry

Global proteomics (“Shotgun proteomics”) was performed using the “bottom-up” method. For this purpose, the protein samples were digested with trypsin and then cleaned with Zip-Tip C18 to extract salts. Then, liquid chromatography was performed with nanoUHPLC Easy nLC 1000 (Thermo Scientific brand, model EASY-nLC 1000, Easy-Spray ColumnPepMap RSLC, P/N ES801) (Thermo Scientific) coupled to a mass spectrometer with Orbitrap technology, which allows a separation of the peptides obtained by tryptic digestion of the sample and subsequent identification. Sample ionization was carried out by nanoelectrospray (Thermo Scientific brand, model EASY-SPRAY. Spray voltage: 1.8 kV). The instrument was equipped with an HCD (High Collision Dissociation) cell and an Orbitrap analyzer yielding the identification of peptides simultaneously to their separation by chromatography. The parameters used during the mass spectrometry analysis were based in FullMS followed by ddMS2 (Data Direct Acquisition or DDA) to identify +1 or multiply charged precursor ions in the mass spectrometry data file. Parent mass (MS) and fragment mass (MS/MS) peak ranges were 300-1800 Da (resolution 70000) and 65-2000 Da (resolution 17500), respectively. The analysis of the “Raw data” delivered by the mass spectrometer was performed using the ProteomeDiscoverer version 2.1 search engine, contrasting the raw data against the sequenced genome of *Nesterenkonia* sp. Act20. The following parameters were set for the search: carbamidomethyl (C) on cysteine was set as fixed; variable modifications included asparagine (N) and glutamine (Q) deamidation and methionine (M) oxidation. Only one missed cleavage was allowed; monoisotopic masses were counted; the precursor peptide mass tolerance was set at 10 ppm; fragment mass tolerance was 0.05 Da. Monoisotopic masses were counted. The MS/MS spectra were searched with Proteome Discoverer v2.1 using a 95% confidence interval (CI%) threshold (P < 0.05). The described protocol was carried out at CEQUIBIEM (by its Spanish acronym, “*Centro de Estudios Químicos y Biológicos for Espectrometría de Masa*”), University of Buenos Aires-Argentina.

### Proteomic data processing and statistical analysis

The CEQUIBIEM service performed raw data analysis and identified proteins and their abundance (measured as the normalized area of the signal intensity of each peptide). Further analyses used the software Perseus v.1.6.2.3, Microsoft Excel, R v.3.6.1, and Cytoscape v.3.7.1 (R Core Team, 2016; Shannon et al., 2003; Tyanova et al., 2016).

By using the Perseus software, the first step of the data set processing was the filtering and selection of the proteins that had two valid abundance values in at least one of the treatments with the consequent elimination of proteins with only one valid abundance value in each treatment, in only one treatment, or without valid values in all four treatments. The next step was the log2 transformation of the standardized areas for each protein. Then, because the invalid values (NaN, “Not a number”) are a product of the inefficiency of the MS method to detect low abundance values, these NaN values were “imputed” (i.e. “randomly replaced”) by the minimum detected values of the normal distribution of the whole data set (Lazar et al., 2016) (Width= 0.5, Down shift= 1.8). Next, the abundance averages for each protein were compared between treatments through the T-significance test with p<0.05. The criteria for determining whether a protein was significantly overregulated in one treatment relative to another treatment were: A) The T-test shows a value of significance p<0.05, B) the value of the difference between the log-scale averages of abundance (log2) being above or below the fold change (FC) 1 and -1 (−1<FC_difference_<1), respectively (see note 1 in the supplementary file 2).

### Annotation and bioinformatics analysis

The sequenced genome of *Nesterenkonia* sp. Act20 has been deposited at DDBJ/ENA/GenBank under the accession JADPQH000000000. The functional analysis of the identified proteins was performed using tools from the “KEGG” (Kyoto Encyclopedia of Genes and Genomes) and String databases (Szklarczyk et al., 2017) and exhaustive text mining.

## RESULTS

### UV-resistance profile of *Nesterenkonia* sp. strains

Act20 and DSM 15474 cultures were assayed on agar media using the drop plate method to qualitative compare the UV-resistance profile of both strains to artificial UV-B radiation and their performance after recovery treatments in light or dark conditions. To check whether an acclimatization process improves cell viability, parallel experiments were performed by exposing two sets of cultured agar plates to equivalent irradiation doses by continuous or discontinuous (pulses) UV-B irradiation (i.e., lethal doses and acclimatization tests, respectively) (Fig. 1).

**Figure 1.**
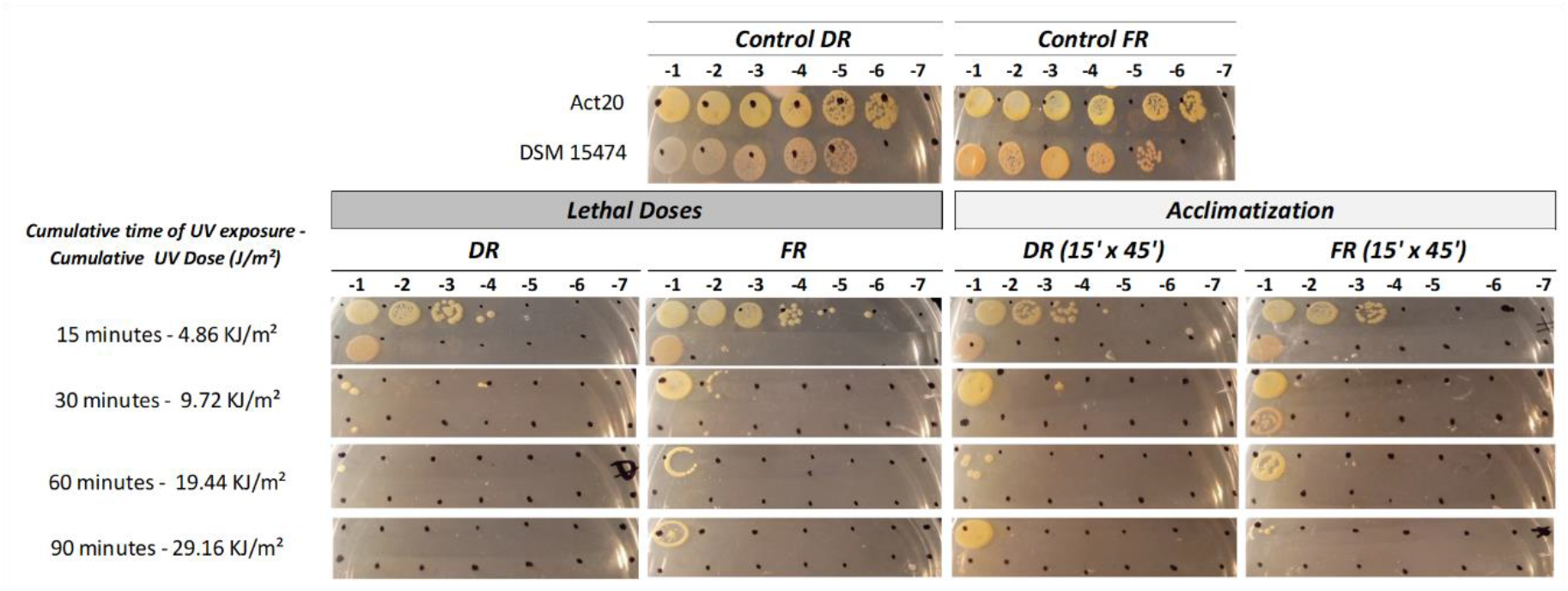
Effects of lethal doses and acclimatization test and dark or white light recovery treatments, respectively, on the cell viability assay of Act20 (top row) and *N. halotolerans* DSM15474 (low row) cultures. DR: cells incubated in the dark after each treatment; PR: cells incubated in PAR light after each treatment.

Act20 and DSM 15474 cultures treated under control conditions showed optimal growth without any observable differences when incubated in the dark (DR) or under light (FR). In contrast, UV-B irradiation limits the growth of both strains, but being much more evident for DSM 15474. The minimal UV-B dose set in our assay reduced the cell viability of Act20 by two or three orders of magnitude depending on the treatment, without any significant notorious influence in recovery treatments. DSM 15474 was inhibited entirely with doses above 9.72 kJ/m² in all treatments, whereas Act20 tolerated higher UV-B doses than *N. halotolerans*, confirming the high UV-resistance profile of the Puna actinobacterium (Alonso-Reyes, Farias, et al., 2020; Alonso-Reyes, Galván, et al., 2020; Portero et al., 2019; Rasuk et al., 2017). Interestingly, the viability of Act20 was proportionally reduced in agreement to the cumulative irradiation when this strain was exposed to continuous UV-B pulses and allowed to recover in darkness.

In contrast, cells survived even at the most intense dose when continuous exposure was followed by photorecovery, and also after treatments using discontinuous UV-B irradiation. Thus, Act20 cultures exposed to UV-B continuous irradiation and recovered on light or exposed to discontinuous pulses and recovered either under light or darkness were significantly impaired by UV-B radiation but remain viable until 29.16 kJ/m² doses. These results indicated that acclimatization seems to improve the adaptive response allowing Act20 to endure high UV intensity. It can also be concluded from these experiments that PAR light has a subtle but beneficial effect over its cellular viability.

### DNA repair ability of *Nesterenkonia* sp. strains

The level of DNA damage upon UV-B exposure and its repair was assessed in Act20 and DSM 15474 by HPLC-ESI-MS/MS. DNA photoproducts were measured in UV-exposed cells and in cultures subjected to recovery treatments in light (FR) and dark (DR), respectively. DNA photoproducts were also quantified in UV-unexposed samples at both the beginning (T0) and the end (Dt) experiment.

The cumulative sum of photoproducts of all categories accumulated within each treatment yields a picture of the damage/repair balance on DNA in both strains under different experimental conditions (fig. 2A). The maximum accumulation of photoproducts was found in DSM 15474 cultures exposed to the UV treatment that reaches a mean cumulative concentration of 1228 photoproducts per million bases (bpm). Surprisingly, Act20 cultures accumulate a lower quantity than DSM 15474 under the same experimental conditions. Thus, if the value found in DSM 15474 UV-exposed cultures is 100% of photoproducts able to be generated by our experimental conditions, then Act20 UV-exposed cultures accumulate about 45% fewer photoproducts ( cf. 671 bpm) than DSM 15474 under the same conditions. A similar pattern was also detected under DR and FR treatments, in which Act20 accumulates ca. 627 and ca. 515 photoproducts per million bases, ≥29% fewer photoproducts than DSM 15474 DR- and FR-exposed cultures, respectively (fig. 2A).

**Figure 2.**
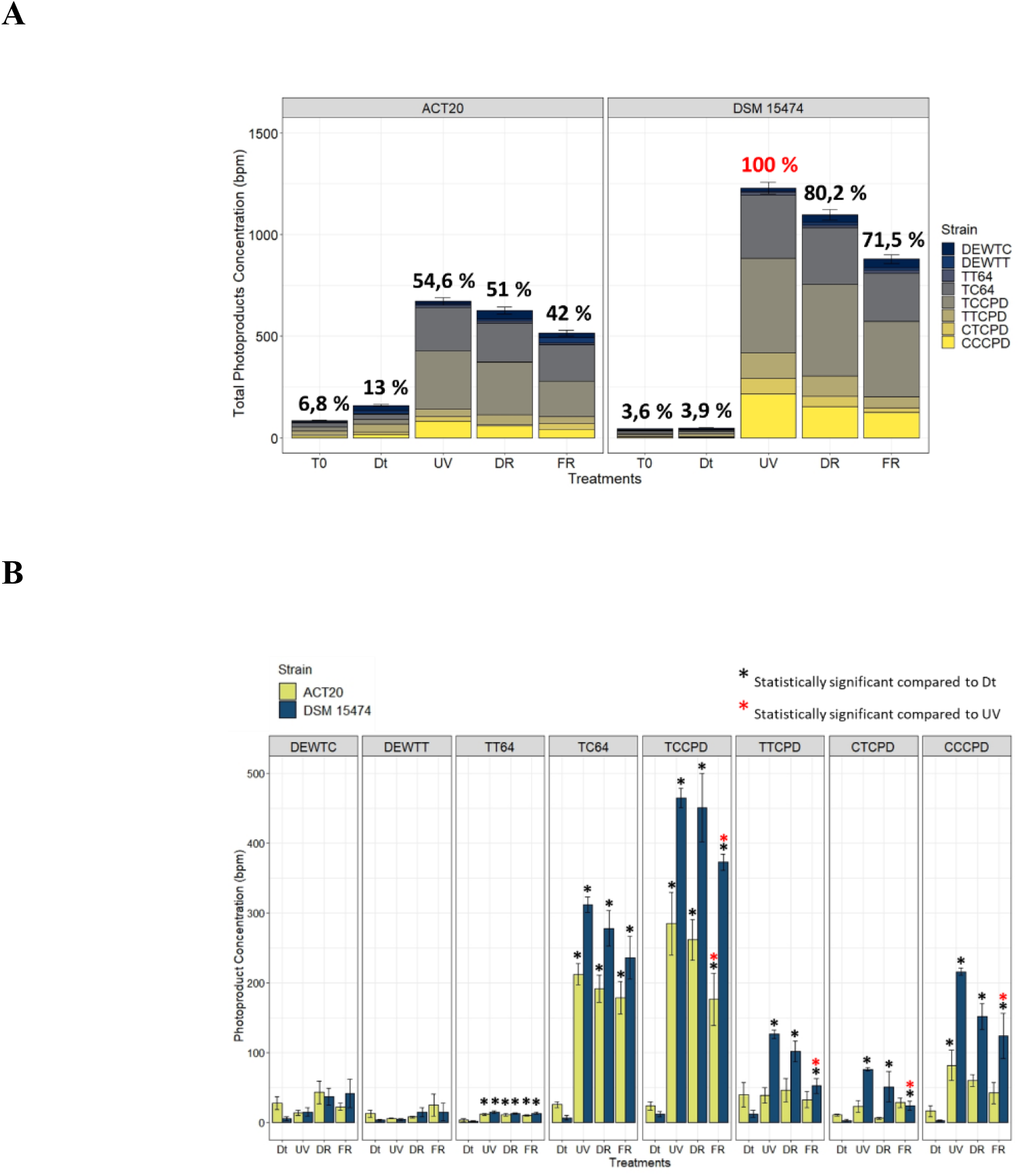
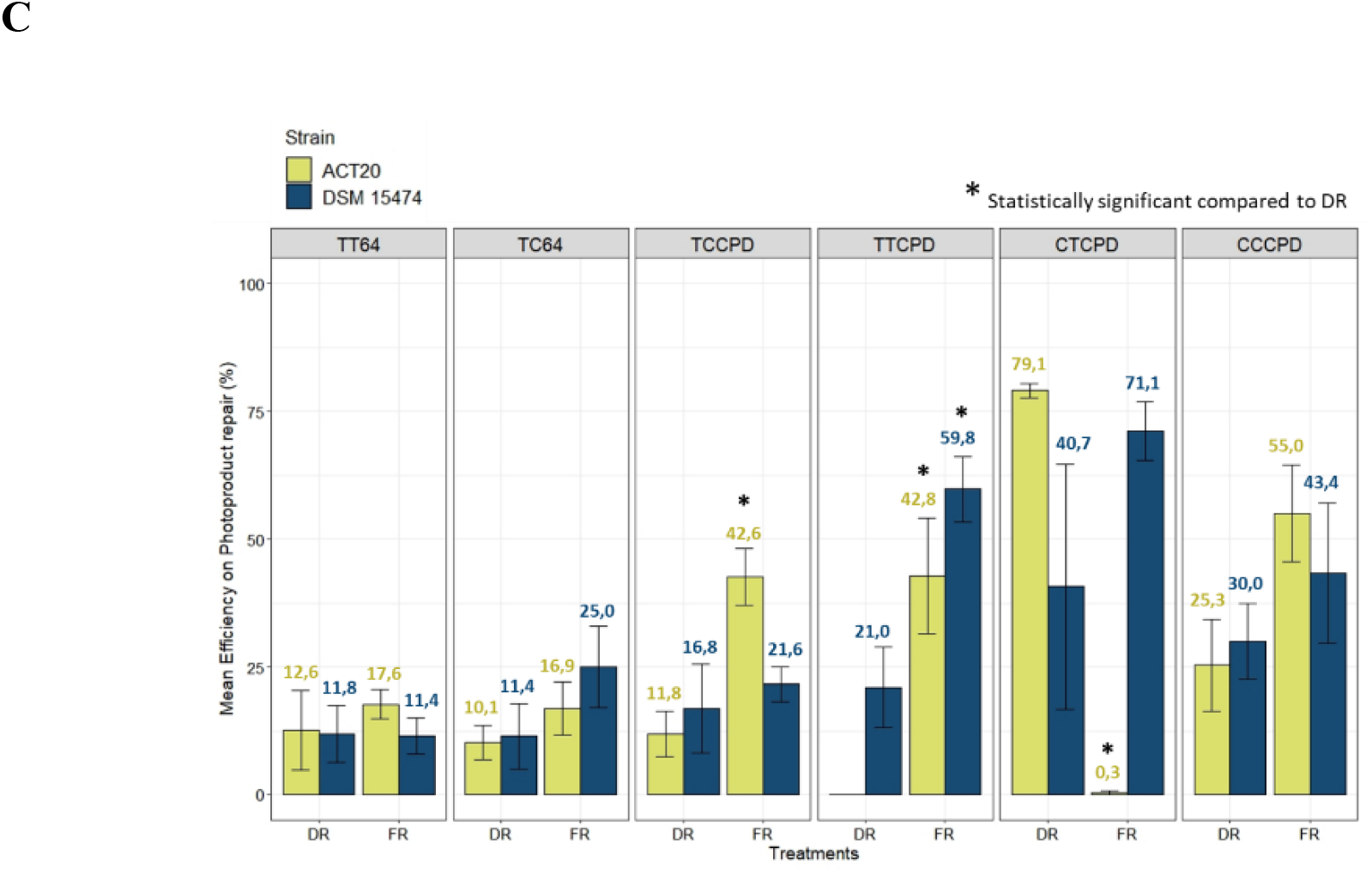
**A)** Stacked barplot showing the mean total concentration (as bases per million (bpm)) of photoproducts generated by each treatment and the contribution of each photoproduct category. Error bars means the standard deviation of the cumulative sum of photoproducts within each treatment. The 100% of photoproducts (in red) able to be produced under our experimental conditions was attributed to the condition with the highest photoproduct concentration found in the UV-treated cultures of DSM 15474. Relations to the 100% are shown as percentages in black. **B)** Quantification of *cis-syn* cyclobutane pyrimidine dimers (TTCPD, TCCPD, CTCPD and CCCPD), Pyrimidine (6-4) pyrimidone photoproducts (TT64 and TC64), and Dewar isomers (DEWTT and DEWTC) plotted as the mean of their absolute concentration (bpm) achieved in each treatment (Dt, UV, DR and FR). **C)** Comparisons of the mean efficiency of DNA photoproducts repair between DR and FR treatments. *Nesterenkonia* sp. Act20 (yellow bars), *Nesterenkonia halotolerans* DSM 15474 (blue bars). Asterisks indicate statistically significant differences (see legends).

Conversely, T0 and Dt treatments induce lower amounts of photoproducts accumulation than UV, DR, and FR treatments in both strains. Under T0 and Dt conditions, Act20 accumulates on the average 83 and ca. 160 photoproducts per million bases, equivalent to 12.4 and 23.8% of its maximum concentration peak detected under UV treatment. Also, such accumulations are equivalent to 6.8, and 13% of the maximum concentration detected in DSM 15474 UV-exposed cultures. These results show that each treatment has effects on the DNA damage/repair balance, in which UV-irradiated cultures show an overall increase in DNA damage while photoproducts accumulation in unexposed cultures remains scarce. Also, they show that Act20 intrinsically accumulates a lower amount of photoproducts than DSM 15474 under UV and recovery treatments while there are reductions in photoproducts total accumulation under either recovery conditions in both strains, especially under FR.

As shown in figure 2B, UV irradiation affects the abundance of all photoproducts categories except Dewar isomers. However, there are differences between strains. Compared to Dt treatment, UV significantly increases the accumulation of TT64, TC64, TCCPD, TTCPD, CTCPD and CCCPD photoproducts in DSM 15474, while only induces significant accumulation of TT64, TC64, TCCPD and CCCPD in Act20. Interestingly, when compared with UV conditions, FR induces a significant reduction in many photoproducts categories. While FR significantly decreases the absolute abundances of TCCPD, TTCPD, CTCPD, and CCPD in DSM 15474, the same experimental conditions significantly reduce the concentration of TCCPD in Act20.

The average efficiency for repairing each photoproduct category shows a different aspect of the photoproduct repair system than comparisons between absolute concentrations, as it represents the average percentage of photoproduct repair within each replicate for each photoproduct category. Thus, the mean efficiency to repair TT64, TC64, TCCPD, TTCPD, CTCPD, and CCCPD photoproducts under DR and FR conditions were estimated for both strains (fig. 2C). In general, repair of photoproducts increases under FR treatments, although those differences are not always statistically significant. Thus, Act20 repairs TCCPD and TTCPD significantly better under light recovery conditions (42.6% and 42.8 %) than under dark conditions (11.8% and 0%). Likewise, DMS 15474 repairs TTCPD more efficiently under light recovery conditions (59.8%) than in the dark (21%). Also, both strains improved their repair efficiency for TC64 and CCCPD photoproducts in the light rather than in the dark, although these differences are not statistically significant. Surprisingly, in Act20 CTCPD photoproducts are better repaired under dark recovery while under light this activity is scarce or even not detected. Besides this, the results suggest that the photorecovery treatment improves the photoproducts repair activity. Since photolyases are the only light-driven enzymes capable of repairing damaged nucleotides in DNA that are detected in both Act20 and DSM15474 genomes, we propose that photolyases are responsible for the decrease in photoproducts concentration under FR conditions. Therefore, photolyases may contribute to protecting the integrity of the genome and they constitute active elements of the UV-resistome in both strains.

### Comparative and functional orthology of Act20 proteomic profile

Four experimental conditions, Dt, UV, DR, and FR, were tested to identify the molecular events involved in the adaptive response of Act20 to artificial UV-B and also after photo and dark recovery. In this section, the proteomic profiles identified in each treatment were analyzed based on mass spectrometry detection limits. Thus, those proteins with invalid values (see Methods) in the three biological replicates were considered as not present in the sample. Taking together the number of proteins detected in each treatment, 1597 different proteins were identified that constitute the herein called experimental proteome (Ex.P) covering 59% of the predicted ORFs of Act20 genome (Predicted proteome, Pred.P, = 2689 ORFs). A lower number of proteins were detected in at least one replica of each treatment, i.e., 1476, 1521, 1546, and 1565 proteins in Dt, UV, DR, and FR datasets, respectively (Fig. 3A).

**Figure 3.**
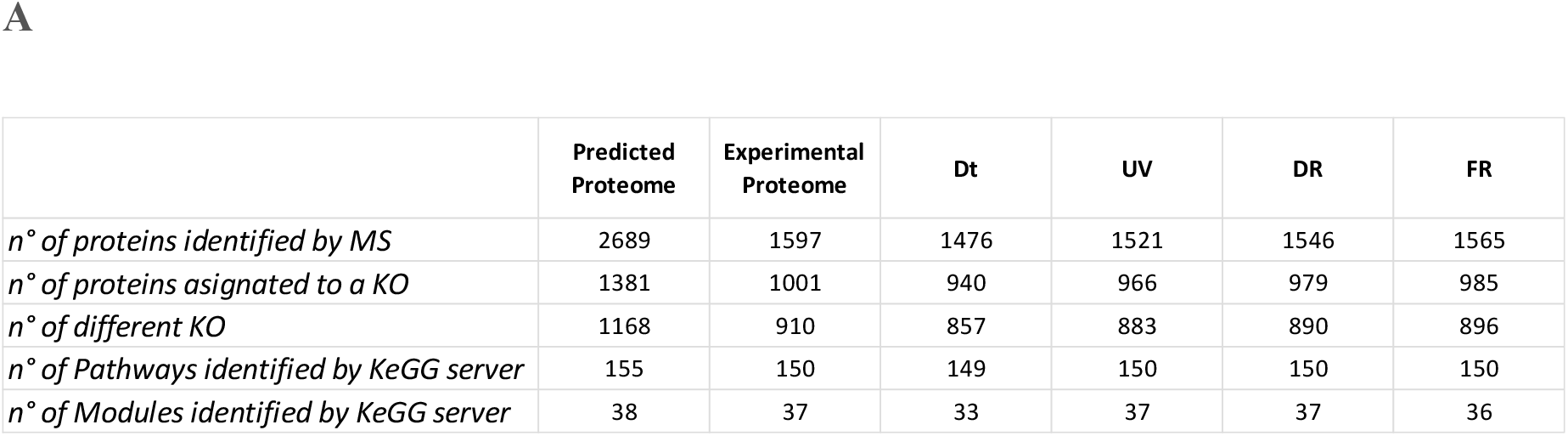
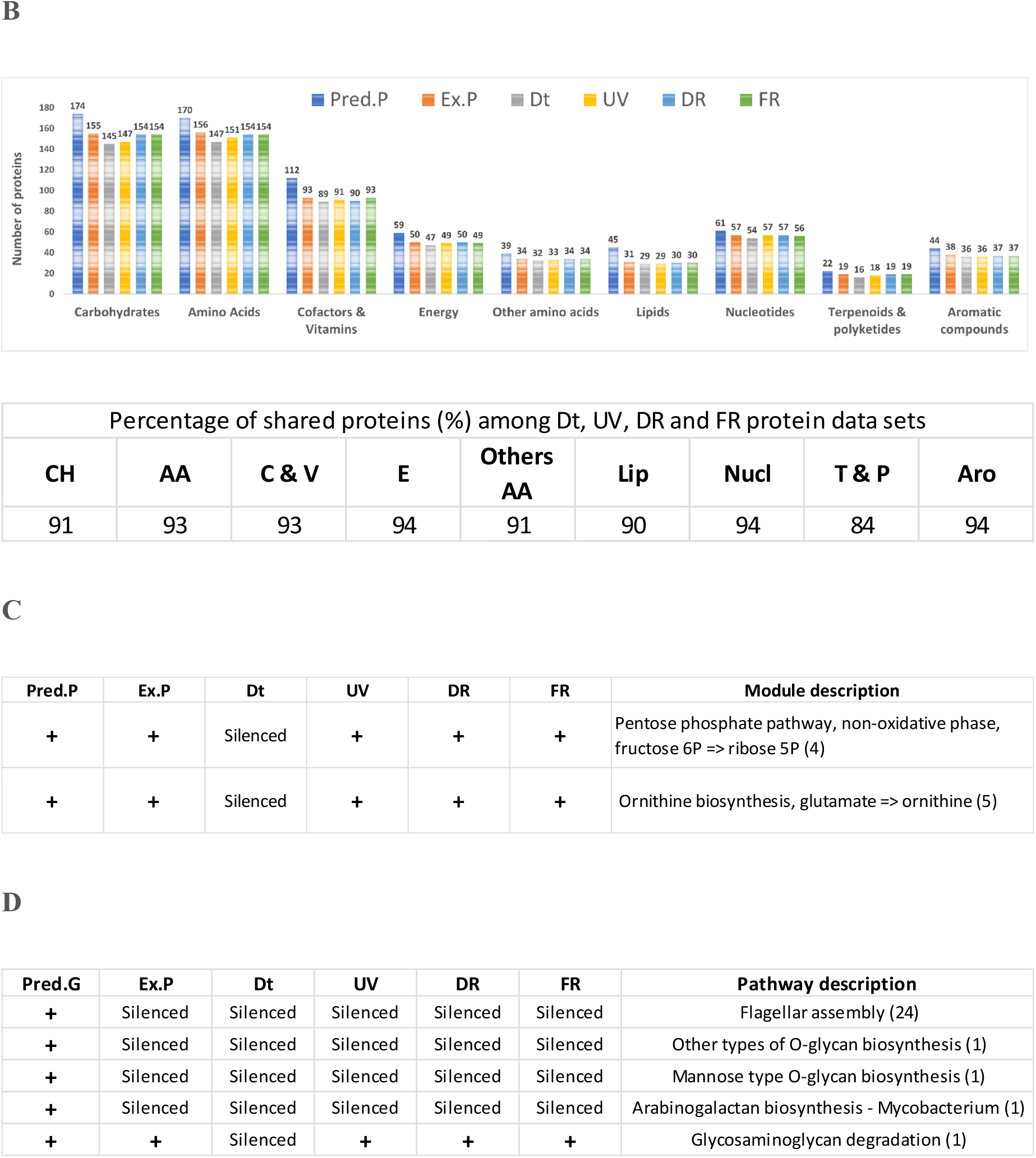
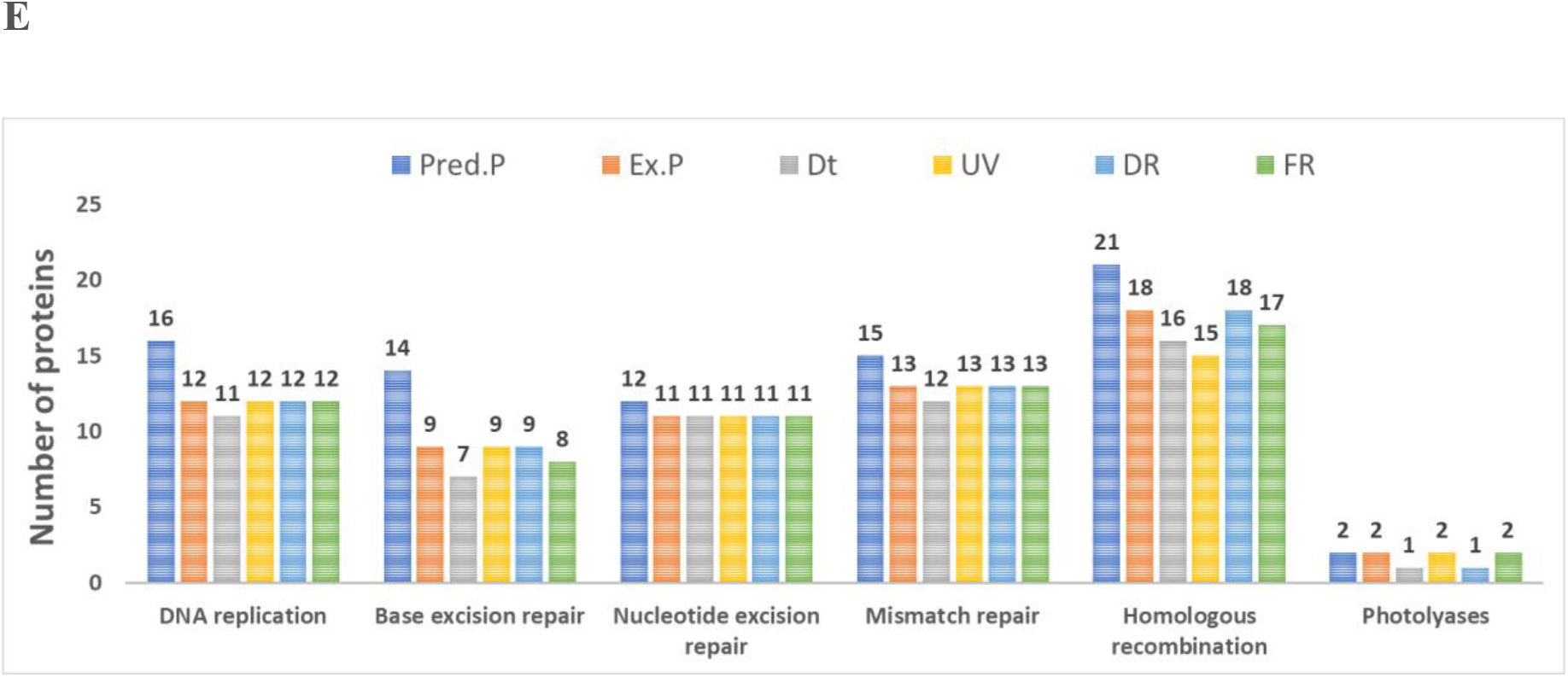
Qualitative analysis of comparative proteomics. **A)** Summary of the number of proteins identified by mass spectrometry and the KEGG database for the individual conditions. **B)** Effect of each condition on the number of proteins involved in the metabolism of the main categories of biomolecules. The table below shows the percentage of proteins shared between the datasets of the experimental treatments for each category of biomolecules, which in turn indicate the percentage of proteins that are replaced by the effect of the treatments. **C)** Summary of functional modules silenced in Dt relative to UV, FR, and DR treatments. **D)** Summary of pathways silenced in the experimental proteome relative to the predicted proteome and those silenced in Dt compared to UV, FR, and DR treatments **E)** Summary of the number of proteins involved in the main categories of DNA repair mechanisms.

In order to reconstruct the metabolic pathways and functional modules contained in each data set of proteins including those contained in the predicted proteome, the experimental proteome and Dt, UV, DR, and FR experimental proteomes, the amino acid sequences of each set of proteins were compared against the KEGG database (Kyoto Encyclopedia of Genes and Genomes). KEGG assigns a code (KO) to each protein according to its function. Several proteins can share the same KO code, but each protein is assigned to only a single KO that reflects its functionality. Thus, KEGG recognized a total of 1381, 1001, 940, 966, 979, and 985 proteins from the predicted proteome, the experimental proteome and Dt, UV, DR, and FR protein data sets, for which 1168, 910, 857, 883, 890 and 896 different protein functionalities were attributed. The lower numbers of different functionalities were observed in Dt data set. In the same way, 155, 150, 149, 150, 150, and 150 metabolic pathways and 38, 37, 33, 37, 37, and 36 functional modules were attributed to Pred.P, Ex.P, Dt, UV, DR, and FR, respectively. The lowest metabolic pathways and functional modules were attributed to the Dt data set (Fig. 3A).

There were no marked alterations induced by Dt, UV, DR or FR on a given metabolism, as the number of proteins detected for each category of biomolecules does not show major variations between datasets and, also, because there was a high percentage of shared proteins among datasets in each particular category (> 90%), indicating a low degree of protein turnover induced by the effect of each experimental condition (Fig. 3B). Thus, the influence of treatments on the metabolism is not dependent on the expression of large groups of proteins, but indicates more the expression and regulation of several specific proteins. Thus, the reconstruction of functional modules using the KEGG server shows that the non-oxidative phase of the pentose phosphate cycle and the biosynthesis of ornithine from glutamate are silenced in the Dt treatment due to the downregulation of specific proteins (i.e., Ribulose-phosphate 3-epimerase (EC:5.1.3.1) (rpe) (see Fig. S1 (Supplemetary File 2)), amino-acid N-acetyltransferase [EC:2.3.1.1] (ArgA), acetylornithine/N-succinyldiaminopimelate aminotransferase [EC:2.6.1.11 2.6.1.17] (ArgD)) (see supplementary file 1), as suggested by the mass spectrometer detection threshold. In contrast, the complete set of proteins involved in these functional modules were detected in UV, DR, and FR protein data sets, respectively, suggesting that such metabolic processes are induced by UV exposure (Fig. 3C). Further, no proteins from the flagellum biosynthesis machinery were detected in the experimental proteome despite being encoded in the predicted proteome, which is consistent with the low motility of the genus *Nesterenkonia*. Also, specific pathways of glycan biosynthesis and metabolism were found to be silenced in the experimental proteome, again despite the fact that these proteins are encoded in the genome of Act20. However, a glycosaminoglycan degradation pathway is silenced in Dt, but not in UV, DR, and FR (Fig. 3D).

As UV-B radiation affects DNA integrity, the number of proteins involved in the primary DNA repair mechanisms was compared showing variation in the number of proteins involved in homologous recombination between treatments. The variation in the photolyase category agrees with the biological function of these enzymes, detecting a second photolyase in UV and FR, but not in Dt or DR in at least one replica (Fig. 3E) (see supplementary file 1).

### Comparison of proteomic profiles among treatments

We selected 1522 out of 1597 proteins from the experimental proteome for a quantitative analysis of the proteome data by filtering and imputing the data set (Lazar et al., 2016) (See Methods). The average abundance values of each protein were compared between treatments data sets by the *t-test*, and the statistical significance of the FC_difference_ was estimated. The effect of artificial UV-B light exposure over the proteomic profile of Act20 was determined mainly by comparing the UV treatment data set against that of Dt treatment (UV-Dt), although FR-UV and DR-UV comparisons were useful to interpret and to complete the model. Likewise, the effect of the photorecovery treatment was determined by comparing the FR data set against Dt and UV (FR-Dt and FR-UV), and, in turn, by comparing each set of FR-upregulated proteins with UV-upregulated and UV-downregulated proteins (from UV-Dt). Also, DR-upregulated proteins taken from DR-Dt and DR-UV comparisons were added to the analysis in order to discriminate between FR-dependent and FR-independent effects on the proteomic profile.

The comparison between UV-Dt data sets revealed fourteen upregulated and twenty-five downregulated proteins (UV challenge) by applying criteria A and B (section Methods) (Figs. 4A, 4B). The intersection analysis between UV-upregulated versus FR-Dt- and DR-Dt-upregulated proteins indicated that there was a set of UV-induced proteins that remains significantly upregulated throughout the recovery treatments (Fig. 4A). As an example, the Frk protein remained upregulated after both recovery treatments, and RecA, DUF4229, and YkoD maintained significantly high levels of abundance during DR and FR, respectively. In addition, this comparison shows that there was a set of 10 proteins exclusively induced by the effect of the UV treatment alone. Similarly, the intersection between the set of UV-upregulated proteins from UV-Dt comparison versus UV-upregulated proteins from FR-UV and DR-UV comparisons indicated that lipA and CdnL were relevant components of Act20 UV-resistome during UV irradiation (Fig. 4C). In contrast, the intersection between the set of UV downregulated proteins (from UV-Dt) versus FR-UV- and DR-UV-upregulated proteins shows the set of proteins that is reduced in abundance due to UV irradiation, but then is restored to their normal levels dependent or independent on FR treatment (Fig. 4B). Thus, RskA, UvrC, adhP, and SPRBCC’ proteins restored their normal levels independently of FR in agreement to criteria A and B, while PC, fdhA, and msmX also restored their normal levels according to criterion A under both recovery treatments, but reached a slightly lower FC_difference_ than that imposed by criterion B in Dt or DR dataset. Despite this, PC, FdhA, and msmX were taken into account because of their close functional link to the FR response model (marked with an asterisk in Fig. 4B and Fig. 6).

**Figure 4.**
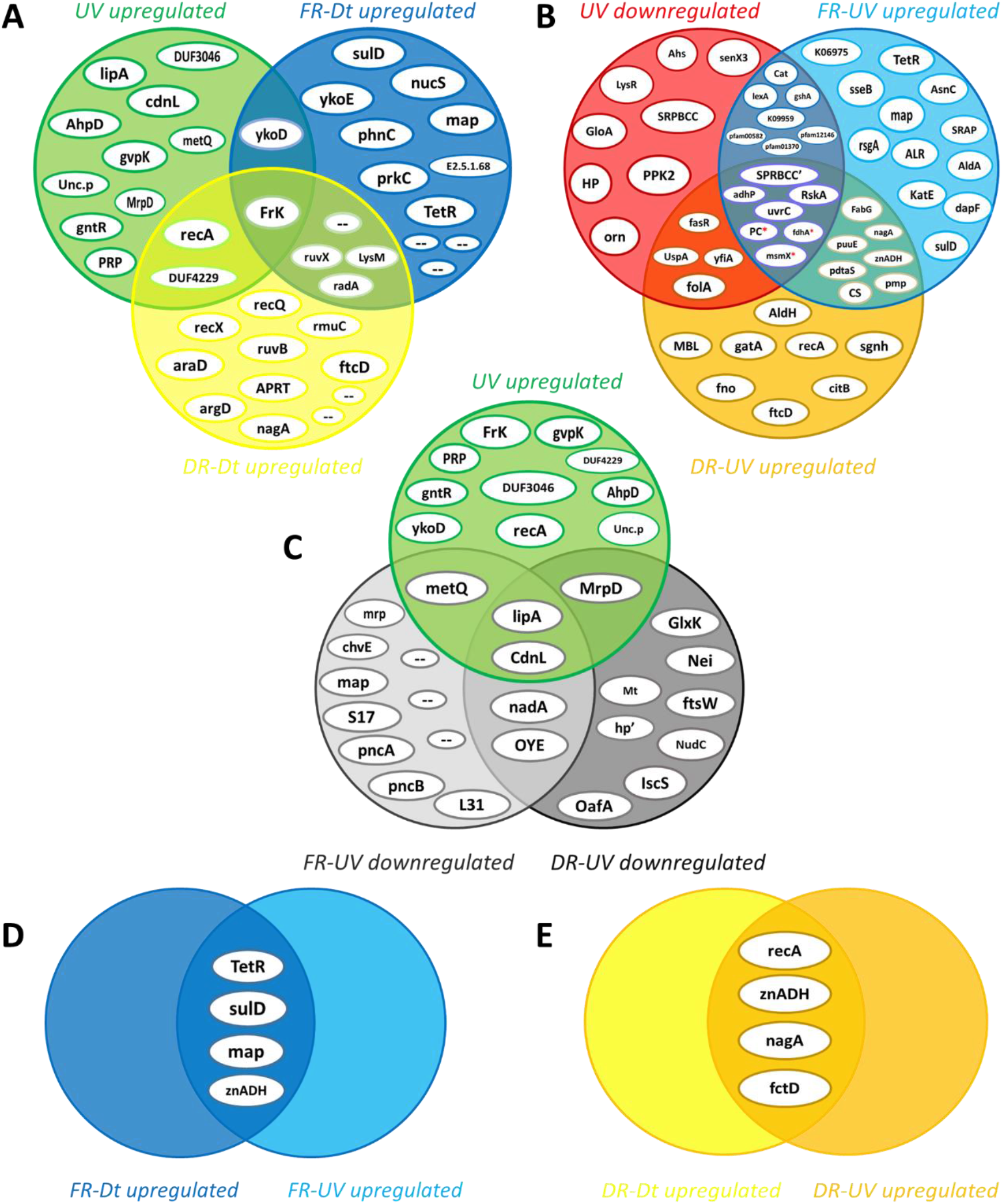
Intersection analysis of upregulated proteins by the effect of each treatment. **A)** A comparison of UV, FR, and DR datasets vs. Dt shows those proteins that remain upregulated in FR and DR after UV irradiation as well as the set of proteins whose abundances were induced during UV exposure. **B)** Comparison of UV-downregulated proteins (from UV-Dt comparison) vs. FR- and DR-upregulated proteins (from FR-UV and DR-UV comparison) indicating the set of proteins that restore their abundance dependently or independently of FR treatment. **C)** Comparison of UV-upregulated proteins from UV-Dt comparison vs. UV-upregulated proteins from FR-UV and DR-UV comparison (FR-UV and DR-UV downregulated, respectively) indicating relevant proteins involved in UV response during irradiation. **D)** Intersection analysis of the sets of FR-upregulated proteins from FR-Dt and FR-UV comparisons indicates relevant proteins involved in FR response due to their high abundance levels values. **E)** Intersection analysis of the sets of DR-upregulated proteins from DR-Dt and DR-UV comparisons.

In addition, lexA, Cat, gshA, K09959, and three uncharacterized proteins related to the universal stress (pfam00582), the nucleoside-diphosphate-sugar epimerases (pfam01370), and the serine aminopeptidases (pfam12146) protein families, recovered their normal levels dependent on FR (Fig. 4B). The abundance levels of the bifunctional folate synthesis protein SulD, the transcriptional regulator of the TetR family (TetR), and the methionine amino peptidase (map) proteins were exclusively increased by FR, while the level of the zinc-dependent aldehyde dehydrogenase (znADH) was increased in both recovery treatments (Fig. 4D, E). SulD, TetR, map and znADH reached abundance values that significantly exceed those achieved in Dt and UV datasets suggesting that they might be central elements of cell recovery under FR (Fig. 4D).

Altogether, these results indicate that UV exposure triggered the upregulation of fourteen main proteins while the photorecovery treatment restored basal levels of critical proteins linked to cell fitness recovery. These results also suggest that once the UV challenge has come to an end, there is a return of the proteomic profile induced by UV toward the basal abundance levels defined by the control treatment (Dt) independently on FR.

### Molecular response model against artificial UV-B radiation

There was a set of fourteen proteins whose abundance levels were significantly increased during UV irradiation in agreement to criteria A and B. In order to build a model of the molecular events involved in the response against UV, each protein was functionally linked to a particular cellular process, when possible. To do this, both the functions and biological implications of each protein were analyzed through extensive literature mining of homologous proteins using cured databases. In addition, to gain more evidences of the molecular events in which the proteins are involved, functionally related proteins (from UV-Dt comparison) with significant FC_difference_ but below the imposed threshold were analyzed and taken into account for the model. Also, significantly UV upregulated proteins from FR-UV and DR-UV comparisons were added (Fig. 4C). Finally, in order to interpret and complete the model, functionally related proteins encoded in Act20 genome, but showing no significant changes in abundance were also added. Through this analysis it was interpreted that the fourteen co-expressed proteins could be functionally linked to hypothesized UV-resistome subsystems (Fig. 5).

**Figure 5.**
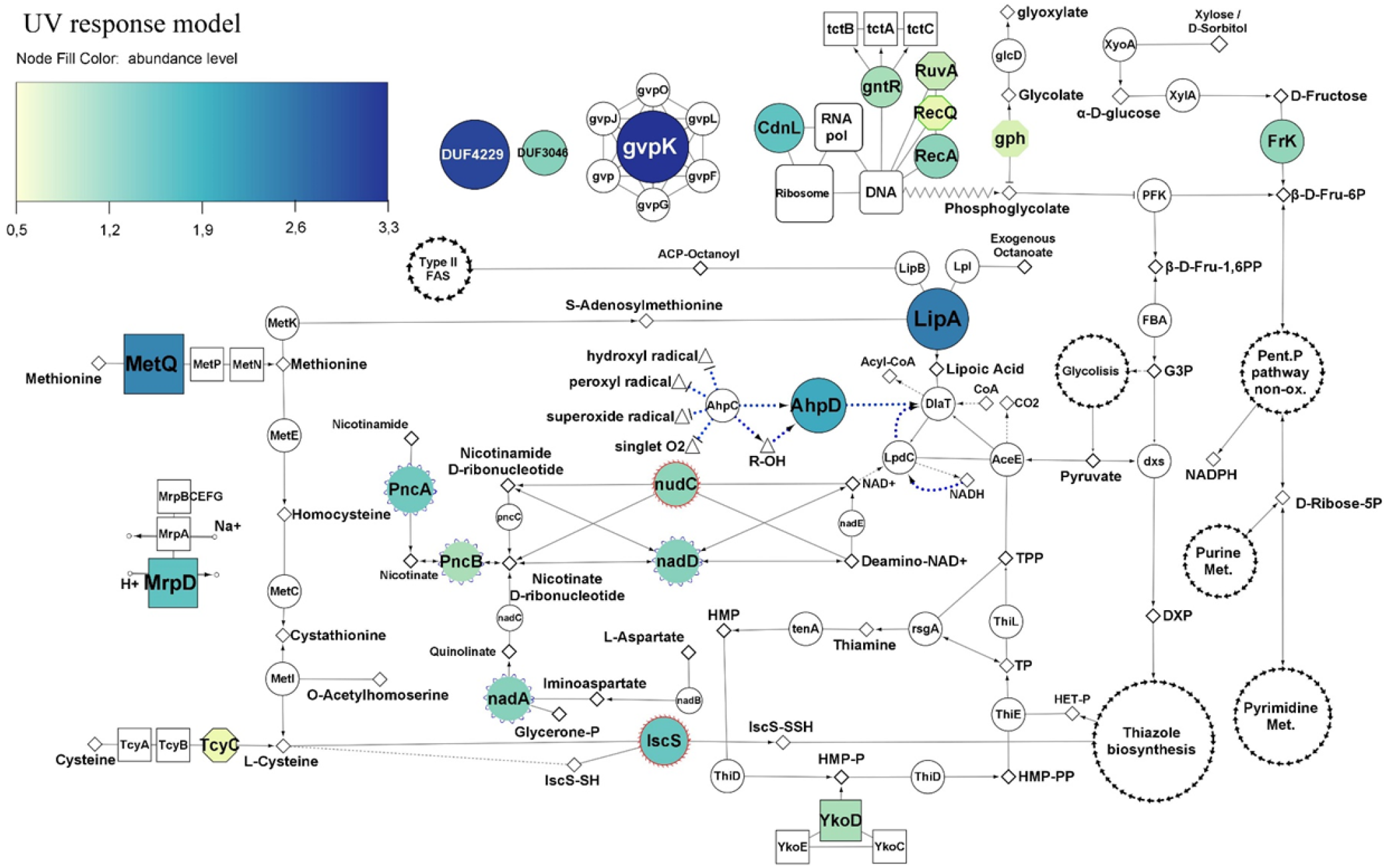
Molecular model of the response to artificial UV-B conditions based on the analysis of the comparative proteomics. The color and size of the nodes indicates the level of abundance of significantly overregulated proteins (see color coding at the top left corner). Color-less nodes indicate functionally associated proteins that show no significant levels of abundance, but are coded in the Act20 genome. The size of these color-less nodes is independent of their level of abundance. Rectangular nodes represent membrane transporters. Nodes with blue scalloped edges (e.g., nadD, in center) and nodes with red edges (e.g., IscS and nudC, center bottom), represent proteins taken from the FR-UV and DR-UV comparisons, respectively. White nodes with separate arrow edges represent cellular processes. Nodes in diamonds, triangles, and small circles represent substrates, chemical compounds, and ions, respectively. Dotted blue arrows indicate the antioxidant activity of the PDH complex.

A first functional module was associated to DNA repair and metabolic resetting. We see that RecA, RecQ, and RuvA proteins are significantly upregulated by the UV effect, suggesting high levels of homologous recombination, an error-free repair system of single strand gaps and double strand breaks on DNA (Khan & Kuzminov, 2012; Kowalczykowski et al., 1994; Kreuzer, 2013; Nickoloff & Hoekstra, 2001; Smith & Wang, 1989). Furthermore, an indirect indicator of DNA damage provoked by UV-induced oxidative stress is the upregulation of the protein phosphoglycolate phosphatase (gph) involved in the dissimilation of the intracellular 2-phosphoglycolate formed during the repair of the 3’-phosphoglycolate ends in the DNA (Pellicer et al., 2003). Another UV upregulated protein was a formamidopyrimidine-DNA glycosylase with endonuclease VIII (nei) activity acts by excising pyrimidines damaged by mutagenic agents (Palmer et al., 1997). CdnL protein (*<u>C</u>ar<u>D N</u>-terminal like*) which belongs to the CarD_CdnL_TRCF family of transcriptional regulators (Bernal-Bernal et al., 2015), was upregulated by UV. This protein has been linked to the global control of gene expression through a RNAP-protein interactions mechanism dependent on σ^A^ factor; its activity is vital and has been well characterized in Mycobacteriales (Garner et al., 2014, 2017; Kaur et al., 2014, 2018; Srivastava et al., 2013; Stallings et al., 2009; Weiss et al., 2012; Zhu et al., 2019) and Myxococcales (Bernal-Bernal et al., 2015; Gallego-García et al., 2014; García-Moreno et al., 2010) under different stress and virulence conditions. It is known that CdnL activates the transcription of rRNA and components of the transcription machinery (Garner et al., 2014, 2017). Although no measurements of ribosomal RNA levels were made in this study, the abundance of a set of ribosomal proteins, whose abundance are proportionally to rRNA expression levels (Garner et al., 2014, 2017), were significantly higher in the UV dataset compared to FR suggesting that ribosomal assembly and metabolic rates (Casati, 2004; Nomura, 1999) may be altered under UV exposure (Fig. S2).

The second functional module was related to the antioxidant activity of the pyruvate dehydrogenase complex (PDH) and to proteins that we interpret as to act systematically to provide the necessary substrates for the synthesis of cofactors required by PDH. In this sense, PDH are enormous protein complexes containing many copies of three proteins named E1 (AceE), E2 (DlaT), and E3 (Lpd*C*) and three cofactors, Lipoic acid (lipoate), Thiamine pyrophosphate (TPP) and NAD+ (Cicchillo et al., 2004; McCarthy & Booker, 2017; Rodionov et al., 2002, 2004, 2008; Spalding & Prigge, 2010). In addition, PDH can be associated with adaptor proteins and act as a powerful antioxidant complex that can efficiently eradicate free radicals generated by UV radiation (Bryk et al., 2002; Jaeger et al., 2004; Lu & Holmgren, 2014; Spalding & Prigge, 2010) (Fig. 5).

The enzyme lipoate synthase (lipA) which is essential for the synthesis of lipoate (Spalding & Prigge, 2010) was upregulated by the UV effect. Also, there were upregulation of proteins linked to the supply of substrate for TPP synthesis from hydroxymethylpyrimidine pyrophosphate (HMP-PP) and hydroxyethyl-thiazole phosphate (HET-P) (Rodionov et al., 2002). Thus, despite Act20 lacks a ThiC enzyme homologue to synthesize HMP-PP from intermediates of the purine biosynthesis pathway, it may introduce exogenous HMP-P into the cell through specific transporters encoded by *YkoCDE* genes and then phosphorylate this compound by Thi*D* to yield HMP-PP. Likewise, Act20 encodes in its genome proteins to synthesize HET-P from derivatives of glycine, cysteine, and the glycolysis metabolism (Fig. S3) (Rodionov et al., 2002). In this model, the enzymatic oxidation of glycine catalyzed by Thi*O* produces iminoglycine (Fig. S3). In turn, the cysteine imported through the Tyc*ABC* membrane transporters, or methionine imported through Met*NPQ* can be transformed into cysteine through the trans-sulphurating pathway and used as a donor of SH groups which are then transferred to the Thi*S* sulfur-carrier-protein by the action of a cysteine desulphurase (Isc*S*) protein yielding ThiS-COSH (Rodionov et al., 2002, 2004) (Fig. 5 and Fig. S3). Simultaneously, intracellular fructose is phosphorylated by Frk (EC 2.7.1.4) and converted into fructose-6P acting as a substrate for glycolysis metabolism, which in turn produces the pyruvate and glyceraldehyde-3-phosphate required by the enzyme 1-deoxy-D-xylulose-5-phosphate synthase (dxs) to yield 1-deoxy-D-xylulose-5-phosphate (DXP) (Rodionov et al., 2002) (Fig. 5 and Fig. S3). Then, iminoglycine, ThiS-COSH, and DXP is condensed by Thi*G* to yield HET-P. Lastly, Thiamin monophosphate is formed by coupling HMP-PP and HET-P. At the next step, thiamin monophosphate is phosphorylated by Thi*L* to form thiamin pyrophosphate (Rodionov et al., 2002). Eventually, TP and TPP can be dephosphorylated by rsgA to yield thiamine that, in turn, can be reused by TenA to produce HMP which can then be reintroduced into the TPP biosynthesis cycle. In this model, YkoD, MetQ, TcyC, IscS, and the fructokinase (EC 2.7.1.4) (FrK) proteins are linked together through the TPP biosynthesis and showed a significant increase of their abundance levels due to the UV effect. Furthermore, in this model, Frk has a dual function, since it can also supply fructose-6P to the non-oxidative phase of the pentose phosphate pathway to yield purine and pyrimidine metabolism intermediates as is evident from the qualitative analysis (Fig. 3C and Fig. S1). In addition, the second functional module includes UV upregulated proteins taken from FR-UV and DR-UV comparisons (Fig. 4C) which are intimately linked to the *de novo* synthesis and regeneration pathways of NAD+ (Rodionov et al., 2008), a cofactor whose function is associated to several reactions, including the functioning of the PDH complex (Spalding & Prigge, 2010). These proteins were PncA, PncB, nadA, nadD and nudC.

As mentioned, the PDH complex can act as an antioxidant by associating DlaT and LpdC proteins with two alkylhydroperoxide reductase peroxyredoxins, AhpC and AhpD. In this system, the oxidized (lipoamide) and reduced (dihydrolipoamide) forms of lipoic acid comprise a redox couple that can effectively extinguish harmful free radicals such as hydroxyl radicals, peroxyl radicals, or superoxide radicals (Bryk et al., 2002; Spalding & Prigge, 2010). Throughout this mechanism, AhpC reduces free radicals and is regenerated by oxidation from AhpD. AhpD is then reduced by oxidation of the dihydrolipoamide linked to DlaT, which in turn is regenerated by LpdC in a NADH-dependent reaction. Therefore, AhpD acts as a bridge between AhpC and the PDH complex suggesting that an increase in its abundance may increase the antioxidant activity of the complex (Fig. 5).

In addition, UV exposure induced a significant increase in the abundance of MrpD protein, which is fundamental in the assembly and functioning of a membrane complex involved in maintaining intracellular homeostasis through extrusion of sodium ions and intrusion of protons (Ito et al., 2017). Also, the abundance levels of gvpK protein (involved in gas vesicle biosynthesis (Pfeifer, 2012)), a GntR transcriptional regulator (probably involved in uptake of citrate and related compounds as suggested from the genomic context analysis (Winnen et al., 2003)), and two uncharacterized protein with DUF4229 and DUF3046 domains, respectively, were significantly increased by UV effect (Fig. 5).

### Effects of the photorecovery treatment on molecular processes after UV-B irradiation

Upon UV-challenge, twenty-five proteins significantly decreased their abundance levels. In order to identify the cellular processes occurring once the UV stimulus has stopped and those stimulated by the photorecovery treatment, the proteins that decreased in abundance under UV irradiation and then restored their normal levels regardless of the recovery condition were of particular interest, as they restored their levels dependent on the effect of FR treatment (Fig. 4B and Fig. 6).

**Figure 6.**
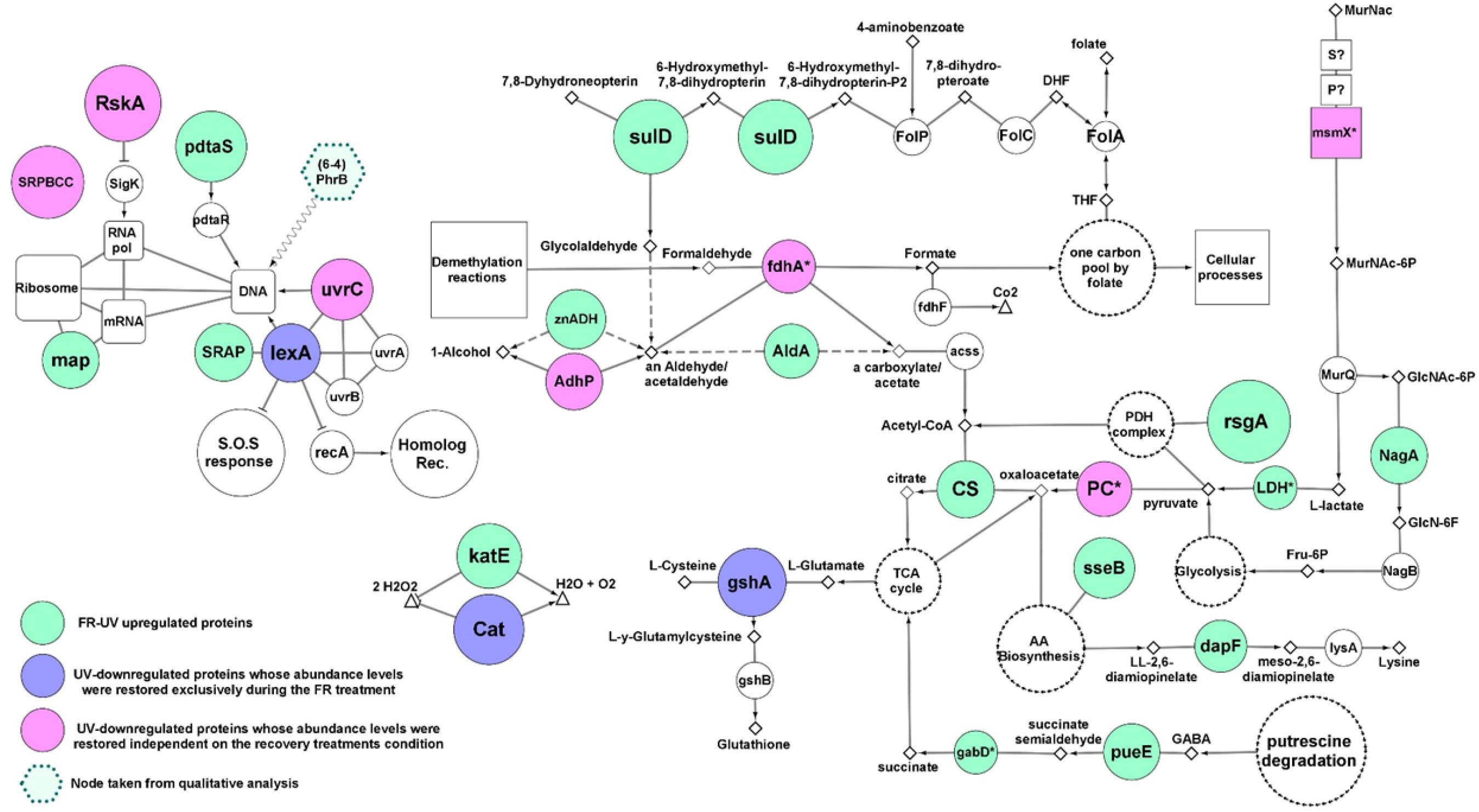
Molecular model of response to FR. Colored nodes indicate proteins identified in experimental proteomics with significantly FC_difference_ (legends in the picture). The size of the nodes indicates the level of abundance. Colorless nodes indicate functionally associated proteins without significantly FC_difference_ but being coded in the Act20 genome. The size of the non-colored nodes is independent of their level of abundance. White nodes with separate arrow heads represent cellular processes.

UV exposure significantly reduced the abundance of the RskA protein, which acts as a repressor of the extracytoplasmic transcriptional factor SigK (ECF19 family) through physical interactions (Huang et al., 2015; Staroń et al., 2009). Then, RskA restored its abundance during FR and DR treatments, suggesting that SigK may coordinate the expression of genes involved in the response against UV-B radiation. UvrC endonuclease involved in nucleotide excision repair of bulky DNA damages such as photoproducts was significantly downregulated under UV, but was restored to its original levels during FR and DR treatments suggesting that the UvrABC base excision repair complex may be involved in the repair of DNA damage caused by UV radiation. Besides, adhP (EC:1.1.1.1), fdhA (EC: 1.2.1.46), Pyruvate carboxylase (PC), and msmX are involved in anaplerotic reactions that supply substrates for the tricarboxylic acid (TCA) cycle through the catabolism of carbohydrates such as alcohols, aldehydes and pyruvate (Arndt et al., 2008; Arndt & Eikmanns, 2007; Gerstmeir et al., 2003; Marçal et al., 2009; Sauer & Eikmanns, 2005; Tanaka et al., 2003) and showed noticeable changes upon UV treatment. Their abundances were significantly decreased under UV exposure and increased later during the recovery treatments, suggesting that energy metabolism and amino acid precursor production from the TCA cycle is truncated by UV irradiation. The involvement of anaplerotic reactions in the recovery response was also supported by the upregulation of the zinc-dependent aldehyde dehydrogenase (znADH), the *Streptomyces aureofaciens* (AAD23400)-like aldehyde dehydrogenase (AldA), Citrate synthase (CS), lactate dehydrogenase (LDH), gabD, and pueE proteins during FR treatment as well as AldH in DR (Jaureguibeitia et al., 2007; Schneider & Reitzer, 2012; Sprušanský et al., 1999) (Fig. 4).

Also, there was a restoration in abundance of both the catalase (Cat) belonging to the manganese cluster family and the glutamate-cysteine ligase gshA involved in thiol biosynthesis. In addition, an upregulation of a second catalase KatE belonging to the heme cluster family was found during FR suggesting a high antioxidant activity under this experimental condition, probably in response to an increase in the amount of free radicals as a consequence of UV irradiation as well as the re-establishment of the metabolic rate (Figs. 4, 6).

Another protein that was found significantly increased in FR was the SOS response repressor lexA. In contrast, the RecA protein remains strongly overregulated in DR indicating that white light may be enhancing the repair of damaged DNA in agreement to the photoproducts repair assay (Fig. 2). Furthermore, the lower abundance values of RecA, radA, and priA proteins in FR treated cultures relative to DR and the upregulation of DNA gyrases and DNA ligases compared to Dt (Fig. S4) are indicative of a temporarily more active state of DNA repair induced by FR. These results support the existence of an efficient photo-induced mechanism of DNA repair possibly involving photolyases as suggested by the qualitative analysis and the photoproducts quantification assay. This finding agrees well with the abundance profile and the biological implications of lexA and RecA proteins (Khan & Kuzminov, 2012; Kreuzer, 2013).

Finally, due to their significantly high abundance values, three unrelated proteins were added to the FR response model, SulD, map, and pdtaS (Fig. 6). SulD is an essential enzyme involved in pterins biosynthesis such as folate, dihydrofolate, and tetrahydrofolate which are basic compounds involved in a wide variety of cellular processes in cell growth and repair (Gaerçon et al., 2006). The methionine amino peptidase (map) participates in the transduction mechanism and protein biosynthesis by removing N-terminal methionine from nascent proteins. Finally, PdtaS is a recently discovered sensor in Actinobacteria whose function and biological implications are currently under discussion. It is proposed that it is involved in the control of amino acid biosynthesis, ribosomal protein levels, and amino acid transfer RNA biosynthesis in a c-di-GMP dependent mechanism (Hariharan et al., 2019; Morth et al., 2005; Preu et al., 2012).

## DISCUSSION

Our results confirm the high UV tolerance profile of *Nesterenkonia* sp. Act20 reported in previous works (Portero et al., 2019) and point out acclimatization and photorecovery as critical strategies to ensure the viability of *Nesterenkonia* sp. cells exposed to high artificial UV-B doses. Both processes represent a tremendous advantage for survival in extreme natural ecosystems such as the High Altitude Andean Lakes as they allow cells to anticipate daily UV fluctuations and, in turn, protect the integrity of cell structure (Albarracín et al., 2012; Portero et al., 2019; Thattai & Van Oudenaarden, 2004; Zenoff, Siñeriz, et al., 2006). A variety of molecular events may accommodate the physiology of the organism and contribute to persist and survive in extreme environments. Through a comparative analytical approach, we have herein identified and integrated the molecular events involved in the adaptive response upon UV-B challenge of Act20 cells. The results showed a functional network of tightly related molecular events involved in the UV-resistome which otherwise Act20 may display in its original ecosystem, the highest UV irradiated environment on Earth.

The comparative proteomic assay revealed that Act20 proteins upregulated by artificial UV-B exposure are part of three proposed UV-resistome subsystems: (i) damage tolerance and oxidative stress response; (ii) energy management and metabolic resetting; and (iii) DNA damage repair. Indeed, homologous recombination was shown to be the primary mechanism of DNA repair as suggested by the high levels of abundance of RecA and other recombinational proteins in UV exposed cultures (Kreuzer, 2013). While homolog recombination acts on single strand gaps and double strand breaks on DNA (secondary lesions), a point mutation (primary lesions) induced by UV irradiation can promote chromosomal fragmentation and single strand gaps when a replication fork advances across a primary lesion which then requires an efficient repair mechanism such as the RecA-dependent processes (Khan & Kuzminov, 2012; Kowalczykowski, 2000; Morimatsu & Kowalczykowski, 2003; Smith & Wang, 1989). Considering the protective mechanism against oxidative stress caused by UV exposure, it seems that the pyruvate dehydrogenase complex is in charge, as there was a significant increase in the abundance of proteins intimately related to its antioxidant activity and the production of the cofactors required for its function, mainly the lipA and AhpD proteins (Spalding & Prigge, 2010). Thus, despite the fact that at first glance ykoD, MetQ, tcyC, IscS, FrK, PncA, PncB, nadA, nadD, and nudC proteins seemed not to be related to the PDH complex, exhaustive literature mining and the review of cured databases led us to the comprehension that these proteins might be linked to the supply of substrates for the production of cofactors such as thiamine pyrophosphate and NAD+ required by this complex (Fig. 5). Equally, methionine imported by the MetQPN transporter is also necessary for the production of S-adenosylmethionine required by the lipoate synthase protein (LipA), further linking also this set of proteins to the PDH complex. Likewise, it is also noteworthy to highlight the importance of the upregulation of Frk (EC:2.7.1.4) (Binet et al., 1998; Caescu et al., 2004; Kelker et al., 1970), as this protein accomplishes a dual function in the UV response model: it can supply substrates for thiamine pyrophosphate biosynthesis through the production of fructose-6P for glycolysis, and it is involved in the production of purine and pyrimidine metabolism intermediates through the non-oxidative phase of the pentose phosphate pathway. In addition, intracellular fructose is the inducer of Frk expression (Binet et al., 1998; Caescu et al., 2004; Kelker et al., 1970). Since exogenous fructose is phosphorylated to fructose-1-phosphate when it enters from outside to inside of the cell through the PTS FruAB II transport system, and that byproduct is neither a substrate or an inducer of Frk (ec 2.7.1.4) (Binet et al., 1998; Caescu et al., 2004; Kelker et al., 1970), the upregulation of Frk (EC:2.7.1.4) suggests that there is a consumption of intracellular fructose sources such as xylose or D-sorbitol (Fig. 5). It should also be kept in mind that Frk remains abundant during recovery treatments. Thus, we consider that FrK (EC:2.7.1.4) is a key enzyme in the UV response, at least under oligotrophic conditions such as physiological solution and under the system tested here.

The photorecovery treatment has been demonstrated that the cell viability of Act20 cultures is improved upon exposure to UV radiation. The analysis of FR upregulated proteins shows that the abundance of the SOS response repressor lexA is significantly increased in FR-exposed cultures, whereas RecA proteins remains highly abundant in DR-exposed cultures indicating that FR-exposed cultures were in a temporarily more advanced state of DNA repair than DR-cultures, since lexA and RecA proteins are intimately linked through a negative feedback circuit induced by the amount of damaged DNA (Kreuzer, 2013). This suggests the existence of an efficient photo-induced mechanism of DNA repair such as photolyases, as indicated by both the qualitative analysis (in which there appears to be present a second type of photolyase under FR treatment, Fig. 3E) and the photoproducts quantification assay in which there are significant reductions of photoproducts concentration in FR treatments (Fig. 2). Replication events at DNA sites bearing cyclobutane pyrimidine dimers and pyrimidine (6-4) pyrimidone photoproducts can generate double strand breaks and single strand gaps in DNA (Khan & Kuzminov, 2012; Kowalczykowski, 2000; Morimatsu & Kowalczykowski, 2003; Smith & Wang, 1989). This induces the upregulation of RecA and other recombinational proteins, in turn triggering effective cleavage of lexA and thereby reducing its abundance. We reasoned that the stimulation of photolyase activity by the photorecovery treatment allows DNA lesions to be repaired before replication forks pass throughout these lesions, thus, preventing ssDNA gaps formation, chromosomal fragmentation, and RecA overexpression. In turn, decreased RecA abundance levels leads to the accumulation of lexA as occurs under FR conditions (Kowalczykowski et al., 1994; Kreuzer, 2013; Nickoloff & Hoekstra, 2001). Thus, photolyases seem to be critical elements of the UV-resistome of Act20 as they are the only light-driven enzymes detected in its genome capable of repairing damaged nucleotides in DNA. In addition, the substantial restoration of the RskA protein abundance levels after UV exposure and during recovery treatments suggests that the extra-cytoplasmic transcriptional factor sigK may coordinate the expression of stress response genes during UV irradiation (Huang et al., 2015; Staroń et al., 2009).

Finally, it is worth noting the remarkable similarity of Act20 molecular UV response mechanisms with those described for common nosocomial strains of the mycobacterial order in response to stress conditions such as the participation of pyruvate dehydrogenase complex through lipA and AhpD proteins, the increased levels of CdnL and the involvement of the RskA-sigK molecular system. Act20 shows also a remarkable similarity to proteins involved in the anaplerotic reaction of the TCA cycle machinery for the catabolism of aldehydes and alcohols compared to the industrial strain *Corynebacterium glutamicum.* Besides, the critical involvement of lipA protein in the UV irradiation response of Act20 suggests greater levels of lipoic acid production, probably due to an increased abundance of free radicals. This supports the idea that lipoate supplementation strategies can contribute to combat oxidative stress. These notes are of utmost important in the context of the current interest in finding strains capable of carrying out biotechnological processes under extreme conditions. Thus, the results from this work aid to highlight the value of microbial communities from th extreme environments such as the Central Andes that need urgent protection from irrational exploitation of their natural environments and ongoing in-depth analysis of their specialized metabolisms.

## REFERENCES

1. Aertsen, A., & Michiels, C. W. (2004). Stress and how bacteria cope with death and survival. Critical Reviews in Microbiology, 30(4), 263–273. https://doi.org/10.1080/10408410490884757

2. Albarracin, V. H., Dib, J. R., Ordoñez, O. F., & Farías, M. E. (2011). A harsh life to indigenous proteobacteria at the andeanmountains: Microbial diversity and resistance mechanisms towards extreme conditions. In M. L. Sezenna (Ed.), Proteobacteria: Phylogeny, Metabolic Diversity and Ecological Effects (pp. 91–130).

3. Albarracín, V. H., Gärtner, W., & Farías, M. E. (2009). UV Resistance and Photoreactivation of Extremophiles from High-Altitude Andean Lakes. 1–20.

4. Albarracín, V. H., Gärtner, W., & Farías, M. E. (2016). Forged Under the Sun : Life and Art of Extremophiles from Andean Lakes. Photochemistry and Photobiology, 92, 14–28. https://doi.org/10.1111/php.12555

5. Albarracín, V. H., Kraiselburd, I., Bamann, C., Wood, P. G., Bamberg, E., Farias, M. E., & G??rtner, W. (2016). Functional green-tuned proteorhodopsin from modern stromatolites. PLoS ONE, 11(5), 1–18. https://doi.org/10.1371/journal.pone.0154962

6. Albarracin, V. H., Kurth, D., Ordoñez, O. F., Belfiore, C., Luccini, E., Salum, G. M., Piacentini, R. D., & Farías, M. E. (2015). High-up: A remote reservoir of microbial extremophiles in central Andean Wetlands. Frontiers in Microbiology, 6(DEC). https://doi.org/10.3389/fmicb.2015.01404

7. Albarracín, V. H., Simon, J., Pathak, G. P., Valle, L., Douki, T., Cadet, J., Borsarelli, C., Farias, M. E., Gärtner, W., & Bossmann, S. (2012). First characterisation of a CPD-class I photolyase from a UV-resistant extremophile isolated from High-Altitude Andean Lakes †. Photochemical & Photobiological Sciences, 1251–1258. https://doi.org/10.1039/c2pp05417e

8. Alonso-Reyes, D., Farias, M., & Albarracín, V. H. (2020). Uncovering cryptochrome/photolyase gene diversity in aquatic microbiomes exposed to diverse UV-B regimes. Aquatic Microbial Ecology, 85(4107), 141–154. https://doi.org/10.3354/ame01947

9. Alonso-Reyes, D., Galván, F. S., Portero, L. R., Alvarado, N., Farías, M. E., Vazquez, M. P., & Albarracín, V. H. (2020). Genomic insights of an andean multi-resistant soil actinobacterium of biotechnological interest. 4107.

10. Arndt, A., Auchter, M., Ishige, T., Wendisch, V. F., & Eikmanns, B. J. (2008). Ethanol catabolism in Corynebacterium glutamicum. Journal of Molecular Microbiology and Biotechnology, 15(4), 222–233. https://doi.org/10.1159/000107370

11. Arndt, A., & Eikmanns, B. J. (2007). The alcohol dehydrogenase gene adhA in Corynebacterium glutamicum is subject to carbon catabolite repression. Journal of Bacteriology, 189(20), 7408–7416. https://doi.org/10.1128/JB.00791-07

12. Belfiore, C., Ordoñez, O. F., & Farías, M. E. (2013). Proteomic approach of adaptive response to arsenic stress in Exiguobacterium sp. S17, an extremophile strain isolated from a high-altitude Andean Lake stromatolite. Extremophiles, 17(3), 421–431. https://doi.org/10.1007/s00792-013-0523-y

13. Bernal-Bernal, D., Gallego-García, A., García-Martínez, G., García-Heras, F., Jiménez, M. A., Padmanabhan, S., & Elías-Arnanz, M. (2015). Structure-function dissection of Myxococcus xanthus CarD N-terminal domain, a defining member of the CarD-CdnL-TRCF family of RNA polymerase interacting proteins. PLoS ONE, 10(3), 1–18. https://doi.org/10.1371/journal.pone.0121322

14. Binet, M. R. B., Rager, M. N., & Bouvet, O. M. M. (1998). Fructose and mannose metabolism in Aeromonas hydrophila: Identification of transport systems and catabolic pathways. Microbiology, 144(4), 1113–1121. https://doi.org/10.1099/00221287-144-4-1113

15. Bryk, R., Lima, C. D., Erdjument-Bromage, H., Tempst, P., & Nathan, C. (2002). Metabolic enzymes of mycobacteria linked to antioxidant defense by a thioredoxin-like protein. Science, 295(5557), 1073–1077. https://doi.org/10.1126/science.1067798

16. Cabrol, N. A., Grin, E. A., Bebout, L., Chong, G., Demergasso, C., Fleming, E., Gaete, V., Gibson, J., Häder, D.-P., Mack, J., Minkley, E., Pinto, E., Rose, K., Ukstins Peate, I., Tambley, C., Williamson, C., & Wynne, J. J. (2009). High lakes project - impact of climate variability and high uv flux on lake habitat: implications for early mars and present-day earth. lpi, 1141.

17. Cabrol, N. A., Grin, E. A., Hock, A., Kiss, A., Borics, G., Kiss, K., Acs, E., Kovacs, G., Chong, G., Demergasso, C., Sivila, R., Ortega Casamayor, E., Zambrana, J., Liberman, M., Sunagua Coro, M., Escudero, L., Tambley, C., Gaete, V., Morris, R. L., … Hovde, G. (2004). Investigating the impact of uv radiation on high-altitude shallow lake habitats, life diversity, and life survival strategies: clues for mars’ past habitability potential lpi, 1049.

18. Caescu, C. I., Vidal, O., Krzewinski, F., & Artenie, V. (2004). Bifidobacterium longum Requires a Fructokinase (Frk; ATP: D. Journal Of Bacteriology, 186(19), 6515–6525. https://doi.org/10.1128/JB.186.19.6515

19. Casati, P. (2004). Crosslinking of Ribosomal Proteins to RNA in Maize Ribosomes by UV-B and Its Effects on Translation. Plant Physiology, 136(2), 3319–3332. https://doi.org/10.1104/pp.104.047043

20. Cicchillo, R. M., Iwig, D. F., Jones, A. D., Nesbitt, N. M., Baleanu-Gogonea, C., Souder, M. G., Tu, L., & Booker, S. J. (2004). Lipoyl synthase requires two equivalents of S-adenosyl-L-methionine to synthesize one equivalent of lipoic acid. Biochemistry, 43(21), 6378–6386. https://doi.org/10.1021/bi049528x

21. Dávila Costa, J. S., Herrero, O. M., Alvarez, H. M., & Leichert, L. (2015). Label-free and redox proteomic analyses of the triacylglycerol-accumulating Rhodococcus jostii RHA1. Microbiology (United Kingdom), 161(3), 593–610. https://doi.org/10.1099/mic.0.000028

22. Dib, J., Weiss, A., Neumann, A., Ordoñez, O., Estévez, M. C., & Farías, M. E. (2009). Isolation of bacteria from remote high altitude Andean lakes able to grow in the presence of antibiotics. Recent Patents on Anti-Infective Drug Discovery, 4(1), 66–76. https://doi.org/10.2174/157489109787236300

23. Douki, T., & Cadet, J. (2001). Individual determination of the yield of the main UV-induced dimeric pyrimidine photoproducts in DNA suggests a high mutagenicity of CC photolesions. Biochemistry, 40(8), 2495–2501. https://doi.org/10.1021/bi0022543

24. Douki, Thierry. (2013). The variety of UV-induced pyrimidine dimeric photoproducts in DNA as shown by chromatographic quantification methods. Photochemical and Photobiological Sciences, 12(8), 1286–1302. https://doi.org/10.1039/c3pp25451h

25. Escudero, L., Chong, G., Demergasso, C., Farías, M. E., Cabrol, N. A., Grin, E., Minkley, Jr., E., & Yu, Y. (2007). Investigating microbial diversity and UV radiation impact at the high-altitude Lake Aguas Calientes, Chile. Instruments, Methods, and Missions for Astrobiology X, 6694(2007), 66940Z. https://doi.org/10.1117/12.736970

26. Farías, M. E. (2020). Microbial Ecosystems in Central Andes Extreme Environments. In Microbial Ecosystems in Central Andes Extreme Environments. https://doi.org/10.1007/978-3-030-36192-1

27. Flores, M. R., Ordoñez, O. F., Maldonado, M. J., & Farías, M. E. (2009). Isolation of UV-B resistant bacteria from two high altitude Andean lakes (4,400 m) with saline and non saline conditions. Journal of General and Applied Microbiology, 55, 447–458. https://doi.org/10.2323/jgam.55.447

28. Gallego-García, A., Mirassou, Y., García-Moreno, D., Elías-Arnanz, M., Jiménez, M. A., & Padmanabhan, S. (2014). Structural insights into RNA polymerase recognition and essential function of Myxococcus xanthus CdnL. PLoS ONE, 9(10). https://doi.org/10.1371/journal.pone.0108946

29. García-Moreno, D., Abellón-Ruiz, J., García-Heras, F., Murillo, F. J., Padmanabhan, S., & Elías-Arnanz, M. (2010). CdnL, a member of the large CarD-like family of bacterial proteins, is vital for Myxococcus xanthus and differs functionally from the global transcriptional regulator CarD. Nucleic Acids Research, 38(14), 4586–4598. https://doi.org/10.1093/nar/gkq214

30. Garçon, A., Levy, C., & Derrick, J. P. (2006). Crystal Structure of the Bifunctional Dihydroneopterin Aldolase/6-hydroxymethyl-7,8-dihydropterin Pyrophosphokinase from Streptococcus pneumoniae. Journal of Molecular Biology, 360(3), 644–653. https://doi.org/10.1016/j.jmb.2006.05.038

31. Garner, A. L., Rammohan, J., Huynh, J. P., Onder, L. M., Chen, J., Bae, B., Jensen, D., Weiss, L. A., Manzano, A. R., Darst, S. A., Campbell, E. A., Nickels, B. E., Galburt, E. A., & Stallings, C. L. (2017). Effects of increasing the affinity of CarD for RNA polymerase on Mycobacterium tuberculosis growth, rRNA transcription, and virulence. Journal of Bacteriology, 199(4). https://doi.org/10.1128/JB.00698-16

32. Garner, A. L., Weiss, L. A., Manzano, A. R., Galburt, E. A., & Stallings, C. L. (2014). CarD integrates three functional modules to promote efficient transcription, antibiotic tolerance, and pathogenesis in mycobacteria. Molecular Microbiology, 93(4), 682–697. https://doi.org/10.1111/mmi.12681

33. Gerstmeir, R., Wendisch, V. F., Schnicke, S., Ruan, H., Farwick, M., Reinscheid, D., & Eikmanns, B. J. (2003). Acetate metabolism and its regulation in Corynebacterium glutamicum. Journal of Biotechnology, 104(1–3), 99–122. https://doi.org/10.1016/S0168-1656(03)00167-6

34. Hariharan, V. N., Thakur, C., Singh, A., Gopinathan, R., Singh, D. P., Sankhe, G., Chandra, N., Bhatt, A., & Saini, D. K. (2019). The histidine kinase PdtaS is a cyclic di-GMP binding metabolic sensor that controls mycobacterial adaptation to nutrient deprivation. BioRxiv, 615575. https://doi.org/http://dx.doi.org/10.1101/615575.

35. Huang, X., Pinto, D., Fritz, G., & Mascher, T. (2015). Environmental sensing in Actinobacteria: A comprehensive survey on the signaling capacity of this phylum. Journal of Bacteriology, 197(15), 2517–2535. https://doi.org/10.1128/JB.00176-15

36. Ito, M., Morino, M., & Krulwich, T. A. (2017). Mrp antiporters have important roles in diverse bacteria and archaea. Frontiers in Microbiology, 8(NOV), 1–12. https://doi.org/10.3389/fmicb.2017.02325

37. Jaeger, T., Budde, H., Flohé, L., Menge, U., Singh, M., Trujillo, M., & Radi, R. (2004). Multiple thioredoxin-mediated routes to detoxify hydroperoxides in Mycobacterium tuberculosis. Archives of Biochemistry and Biophysics, 423(1), 182–191. https://doi.org/10.1016/j.abb.2003.11.021

38. Jaureguibeitia, A., Saá, L., Llama, M. J., & Serra, J. L. (2007). Purification, characterization and cloning of aldehyde dehydrogenase from Rhodococcus erythropolis UPV-1. Applied Microbiology and Biotechnology, 73(5), 1073–1086. https://doi.org/10.1007/s00253-006-0558-4

39. Kærn, M., Elston, T. C., Blake, W. J., & Collins, J. J. (2005). Stochasticity in gene expression: From theories to phenotypes. Nature Reviews Genetics, 6(6), 451–464. https://doi.org/10.1038/nrg1615

40. Kaur, G., Dutta, D., & Thakur, K. G. (2014). Crystal structure of Mycobacterium tuberculosis CarD, an essential RNA polymerase binding protein, reveals a quasidomain-swapped dimeric structural architecture. Proteins: Structure, Function and Bioinformatics, 82(5), 879–884. https://doi.org/10.1002/prot.24419

41. Kaur, G., Kaundal, S., Kapoor, S., Grimes, J. M., Huiskonen, J. T., & Thakur, K. G. (2018). Mycobacterium tuberculosis CarD, an essential global transcriptional regulator forms amyloid-like fibrils. Scientific Reports, 8(1), 1–13. https://doi.org/10.1038/s41598-018-28290-4

42. Kelker, N. E., Hanson, T. E., & Anderson, R. L. (1970). Alternate Pathways of D-Fructose in Aerobacter aerogenes. The Journal of Biological Chemistry, 245(8), 2060–2065. https://doi.org/https://doi.org/10.1016/S0021-9258(18)63206-5

43. Khan, S. R., & Kuzminov, A. (2012). Replication forks stalled at ultraviolet lesions are rescued via RecA and RuvABC protein-catalyzed disintegration in Escherichia coli. Journal of Biological Chemistry, 287(9), 6250–6265. https://doi.org/10.1074/jbc.M111.322990

44. Kowalczykowski, S. C. (2000). Initiation of genetic recombination and recombination-dependent replication. Trends in Biochemical Sciences, 25(4), 156–165. https://doi.org/10.1016/S0968-0004(00)01569-3

45. Kowalczykowski, S. C., Dixon, D. A., Eggleston, A. K., Lauder, S. D., & Rehrauer, W. M. (1994). Biochemistry of homologous recombination in Escherichia coli. Microbiological Reviews, 58(3), 401–465. https://doi.org/10.1177/0898264309358764

46. Kreuzer, K. N. (2013). DNA Damage Responses in Prokaryotes : Replication Forks. Cold Spring Harb Perspect Biol, 1–23.

47. Kurth, D., Amadio, A., Ordoñez, O. F., Albarracín, V. H., Gärtner, W., & Farías, M. E. (2017). Arsenic metabolism in high altitude modern stromatolites revealed by metagenomic analysis /631/326/47/4112/631/326/2565/2142 /45/23 /45/22 article. Scientific Reports, 7(1), 0–16. https://doi.org/10.1038/s41598-017-00896-0

48. Kurth, D., Belfiore, C., Gorriti, M. F., Cortez, N., Farias, M. E., & Albarracín, V. H. (2015). Genomic and proteomic evidences unravel the UV-resistome of the poly-extremophile Acinetobacter sp. Ver3. Frontiers in Microbiology, 6(APR), 1–18. https://doi.org/10.3389/fmicb.2015.00328

49. Lazar, C., Gatto, L., Ferro, M., Bruley, C., & Burger, T. (2016). Accounting for the Multiple Natures of Missing Values in Label-Free Quantitative Proteomics Data Sets to Compare Imputation Strategies. Journal of Proteome Research, 15(4), 1116–1125. https://doi.org/10.1021/acs.jproteome.5b00981

50. Lu, J., & Holmgren, A. (2014). The thioredoxin antioxidant system. Free Radical Biology and Medicine, 66, 75–87. https://doi.org/10.1016/j.freeradbiomed.2013.07.036

51. Marçal, D., Rêgo, A. T., Carrondo, M. A., & Enguita, F. J. (2009). 1,3-Propanediol dehydrogenase from Klebsiella pneumoniae: Decameric quaternary structure and possible subunit cooperativity. Journal of Bacteriology, 191(4), 1143–1151. https://doi.org/10.1128/JB.01077-08

52. McCarthy, E. L., & Booker, S. J. (2017). Destruction and reformation of an iron-sulfur cluster during catalysis by lipoyl synthase. Science, 358(6361), 373–377. https://doi.org/10.1126/science.aan4574

53. Morimatsu, K., & Kowalczykowski, S. C. (2003). RecFOR proteins load RecA protein onto gapped DNA to accelerate DNA strand exchange: A universal step of recombinational repair. Molecular Cell, 11(5), 1337–1347. https://doi.org/10.1016/S1097-2765(03)00188-6

54. Morth, J. P., Gosmann, S., Nowak, E., & Tucker, P. A. (2005). A novel two-component system found in Mycobacterium tuberculosis. FEBS Letters, 579(19), 4145–4148. https://doi.org/10.1016/j.febslet.2005.06.043

55. Nickoloff, J. A., & Hoekstra, M. F. (2001). DNA Damage and Repair. In Springer Science & Business Media: Vol. Volume III (Issue Advances from phage to humans). https://doi.org/10.1038/nature01408

56. Nomura, M. (1999). Regulation of Ribosome Biosynthesis in Escherichia coli and Saccharomyces cerevisiae : Diversity and Common GUEST COMMENTARY Regulation of Ribosome Biosynthesis in Escherichia coli and Saccharomyces cerevisiae : Diversity and Common Principles. Journal of Bacteriology, 181(22), 6857–6864.

57. Ordoñez, Omar F., Flores, M. R., Dib, J. R., Paz, A., & Farías, M. E. (2009). Extremophile culture collection from Andean lakes: Extreme pristine environments that host a wide diversity of microorganisms with tolerance to UV radiation. Microbial Ecology, 58(3), 461–473. https://doi.org/10.1007/s00248-009-9527-7

58. Ordoñez, Omar Federico, Rasuk, M. C., Soria, M. N., Contreras, M., & Farías, M. E. (2018). Haloarchaea from the Andean Puna: Biological Role in the Energy Metabolism of Arsenic. Microbial Ecology, 76(3), 695–705. https://doi.org/10.1007/s00248-018-1159-3

59. Orellana, R., Rojas, C., Seeger, M., Cumsille, A., Macaya, C., Dorochesi, F., Bravo, G., & Valencia, R. (2018). Living at the Frontiers of Life: Extremophiles in Chile and Their Potential for Bioremediation. Frontiers in Microbiology, 9. https://doi.org/10.3389/fmicb.2018.02309

60. Palmer, C. M., Serafini, D. M., & Schellhorn, H. E. (1997). Near ultraviolet radiation (UVA and UVB) causes a formamidopyrimidineglycosylase-dependent increase in G to T transversions. Photochem Photobiol, 65(3), 543–549.

61. Pellicer, M. T., Nun, M. F., & Baldoma, L. (2003). Role of 2-Phosphoglycolate Phosphatase of. Microbiology, 185(19), 5815–5821. https://doi.org/10.1128/JB.185.19.5815

62. Perez, M. F., Kurth, D., Farías, M. E., Soria, M. N., Castillo Villamizar, G. A., Poehlein, A., Daniel, R., & Dib, J. R. (2020). First Report on the Plasmidome From a High-Altitude Lake of the Andean Puna. Frontiers in Microbiology, 11(June), 1–15. https://doi.org/10.3389/fmicb.2020.01343

63. Pérez, V., Hengst, M., Kurte, L., Dorador, C., Jeffrey, W. H., Wattiez, R., Molina, V., & Matallana-Surget, S. (2017a). Bacterial survival under extreme UV radiation: A comparative proteomics study of Rhodobacter sp., isolated from high altitude wetlands in Chile. Frontiers in Microbiology, 8(JUN), 1–16. https://doi.org/10.3389/fmicb.2017.01173

64. Pérez, V., Hengst, M., Kurte, L., Dorador, C., Jeffrey, W. H., Wattiez, R., Molina, V., & Matallana-Surget, S. (2017b). Bacterial survival under extreme UV radiation: A comparative proteomics study of Rhodobacter sp., isolated from high altitude wetlands in Chile. Frontiers in Microbiology, 8(JUN), 1–16. https://doi.org/10.3389/fmicb.2017.01173

65. Pfeifer, F. (2012). Distribution, formation and regulation of gas vesicles. Nature Reviews Microbiology, 10(10), 705–715. https://doi.org/10.1038/nrmicro2834

66. Portero, L. R., Alonso-Reyes, D. G., Zannier, F., Vazquez, M. P., Farías, M. E., Gärtner, W., & Albarracín, V. H. (2019). Photolyases and Cryptochromes in UV-resistant Bacteria from High-altitude Andean Lakes. Photochemistry and Photobiology, 95(1), 315–330. https://doi.org/10.1111/php.13061

67. Preu, J., Panjikar, S., Morth, P., Jaiswal, R., Karunakar, P., & Tucker, P. A. (2012). The sensor region of the ubiquitous cytosolic sensor kinase, PdtaS, contains PAS and GAF domain sensing modules. Journal of Structural Biology, 177(2), 498–505. https://doi.org/10.1016/j.jsb.2011.11.012

68. R Core Team. (2016). R: A language and environment for statistical computing. R Foundation for Statistical Computing. https://www.r-project.org/

69. Ramírez Santos, J., Solís Guzmán, G., & Gómez Eichelmann, M. C. (2001). Regulación genética en la respuesta al estrés calórico en Escherichia coli. Revista Latinoamericana de Microbiologia, 43(1), 51–63.

70. Rascovan, Nicolas, Javier, M., Martín P, V., & María, E. F. (2015). Metagenomic study of red biofilms from Diamante Lake reveals ancient arsenic bioenergetics in haloarchaea Metagenomic study of red biofilms from Diamante Lake reveals ancient arsenic bioenergetics in haloarchaea. International Society for Microbial Ecology, 109(July), 1–11. https://doi.org/10.1038/ismej.2015.109

71. Rascovan, Nicolás, Maldonado, J., Vazquez, M. P., & Eugenia Farías, M. (2016). Metagenomic study of red biofilms from Diamante Lake reveals ancient arsenic bioenergetics in haloarchaea. ISME Journal, 10(2), 299–309. https://doi.org/10.1038/ismej.2015.109

72. Rasuk, M. C., Ferrer, G. M., Kurth, D., Portero, L. R., Farías, M. E., & Albarracín, V. H. (2017). UV-Resistant Actinobacteria from High-Altitude Andean Lakes: Isolation, Characterization and Antagonistic Activities. Photochemistry and Photobiology, 93(3), 865–880. https://doi.org/10.1111/php.12759

73. Rodionov, D. A., De Ingeniis, J., Mancini, C., Cimadamore, F., Zhang, H., Osterman, A. L., & Raffaelli, N. (2008). Transcriptional regulation of NAD metabolism in bacteria: NrtR family of Nudix-related regulators. Nucleic Acids Research, 36(6), 2047–2059. https://doi.org/10.1093/nar/gkn047

74. Rodionov, D. A., Vitreschak, A. G., Mironov, A. A., & Gelfand, M. S. (2002). Comparative genomics of thiamin biosynthesis in procaryotes. New genes and regulatory mechanisms. Journal of Biological Chemistry, 277(50), 48949–48959. https://doi.org/10.1074/jbc.M208965200

75. Rodionov, D. A., Vitreschak, A. G., Mironov, A. A., & Gelfand, M. S. (2004). Comparative genomics of the methionine metabolism in Gram-positive bacteria: A variety of regulatory systems. Nucleic Acids Research, 32(11), 3340–3353. https://doi.org/10.1093/nar/gkh659

76. Saona, L. A., Soria, M., Durán-Toro, V., Wörmer, L., Milucka, J., Castro-Nallar, E., Meneses, C., Contreras, M., & Farías, M. E. (2021). Phosphate-Arsenic Interactions in Halophilic Microorganisms of the Microbial Mat from Laguna Tebenquiche: from the Microenvironment to the Genomes. Microb Ecol. 2021 Jan 2. doi: 10.1007/s00248-020-01673-9. Epub ahead of print. PMID: 33388944. *Microb Ecology 2*. https://doi.org/doi: 10.1007/s00248-020-01673-9

77. Sauer, U., & Eikmanns, B. J. (2005). The PEP-pyruvate-oxaloacetate node as the switch point for carbon flux distribution in bacteria. FEMS Microbiology Reviews, 29(4), 765–794. https://doi.org/10.1016/j.femsre.2004.11.002

78. Schneider, B. L., & Reitzer, L. (2012). Pathway and enzyme redundancy in putrescine catabolism in Escherichia coli. Journal of Bacteriology, 194(15), 4080–4088. https://doi.org/10.1128/JB.05063-11

79. Shannon, P., Markiel, A., Ozier, O., Baliga, N. S., Wang, J. T., Ramage, D., Amin, N., Schwikowski, B., & Ideker, T. (2003). Cytoscape: A Software Environment for Integrated Models of Biomolecular Interaction Networks. Genome Research, 13(11):249.

80. Smith, K. C., & Wang, T. V. (1989). recA-dependent DNA repair processes. BioEssays, 10(1), 12–16. https://doi.org/10.1002/bies.950100104

81. Spalding, M. D., & Prigge, S. T. (2010). Lipoic Acid Metabolism in Microbial Pathogens. Microbiology and Molecular Biology Reviews, 74(2), 200–228. https://doi.org/10.1128/mmbr.00008-10

82. Sprušanský, O., Homérová, D., Ševčíková, B., & Kormanec, J. (1999). Cloning of the putative aldehyde dehydrogenase, aldA, gene from Streptomyces aureofaciens. Folia Microbiologica, 44(5), 491–502. https://doi.org/10.1007/BF02816249

83. Srivastava, D. B., Leon, K., Osmundson, J., Garner, A. L., Weiss, L. A., Westblade, L. F., Glickman, M. S., Landick, R., Darst, S. A., Stallings, C. L., & Campbell, E. A. (2013). Structure and function of CarD, an essential mycobacterial transcription factor. Proceedings of the National Academy of Sciences of the United States of America, 110(31), 12619–12624. https://doi.org/10.1073/pnas.1308270110

84. Stallings, C. L., Stephanou, N. C., Chu, L., Hochschild, A., Nickels, B. E., & Glickman, M. S. (2009). CarD Is an Essential Regulator of rRNA Transcription Required for Mycobacterium tuberculosis Persistence. Cell, 138(1), 146–159. https://doi.org/10.1016/j.cell.2009.04.041

85. Staroń, A., Sofia, H. J., Dietrich, S., Ulrich, L. E., Liesegang, H., & Mascher, T. (2009). The third pillar of bacterial signal transduction: Classification of the extracytoplasmic function (ECF) σ factor protein family. Molecular Microbiology, 74(3), 557–581. https://doi.org/10.1111/j.1365-2958.2009.06870.x

86. Szklarczyk, D., Morris, J. H., Cook, H., Kuhn, M., Wyder, S., Simonovic, M., Santos, A., Doncheva, N. T., Roth, A., Bork, P., Jensen, L. J., & Von Mering, C. (2017). The STRING database in 2017: Quality-controlled protein-protein association networks, made broadly accessible. Nucleic Acids Research, 45(D1), D362–D368. https://doi.org/10.1093/nar/gkw937

87. Tanaka, N., Kusakabe, Y., Ito, K., Yoshimoto, T., & Nakamura, K. T. (2003). Crystal structure of glutathione-independent formaldehyde dehydrogenase. Chemico-Biological Interactions, 143–144, 211–218. https://doi.org/10.1016/S0009-2797(02)00168-0

88. Thattai, M., & Van Oudenaarden, A. (2004). Stochastic gene expression in fluctuating environments. Genetics, 167(1), 523–530. https://doi.org/10.1534/genetics.167.1.523

89. Toneatti, D. M., Albarracín, V. H., Flores, M. R., Polerecky, L., & Farías, M. E. (2017). Stratified bacterial diversity along physico-chemical gradients in high-altitude modern stromatolites. Frontiers in Microbiology, 8(APR), 1–12. https://doi.org/10.3389/fmicb.2017.00646

90. Tyanova, S., Temu, T., Sinitcyn, P., Carlson, A., Hein, M. Y., Geiger, T., Mann, M., & Cox, J. (2016). The Perseus computational platform for comprehensive analysis of (prote)omics data. Nature Methods, 13(9), 731–740. https://doi.org/10.1038/nmeth.3901

91. Vignale, F. A., Lencina, A. I., Stepanenko, T. M., Soria, M. N., Saona, L. A., Kurth, D., Guzmán, D., Foster, J. S., Poiré, D. G., Villafañe, P. G., Albarracín, V. H., Contreras, M., & Farías, M. E. (2021). Lithifying and non-lithifying microbial ecosystems in the wetlands and salt flats of the Central Andes. Microbial Ecology, 1–17. https://doi.org/10.1007/s00248-021-01725-8

92. Weiss, L. A., Harrison, P. G., Nickels, B. E., Glickman, M. S., Campbell, E. A., Darst, S. A., & Stallings, C. L. (2012). Interaction of CarD with RNA Polymerase Mediates Mycobacterium tuberculosis Viability, Rifampin Resistance, and Pathogenesis. Journal of Bacteriology, 194(20), 5621–5631. https://doi.org/10.1128/jb.00879-12

93. Winnen, B., Hvorup, R. N., & Saier, M. H. (2003). The tripartite tricarboxylate transporter (TTT) family. Research in Microbiology, 154(7), 457–465. https://doi.org/10.1016/S0923-2508(03)00126-8

94. Zannier, F., Portero, L. R., Ordoñez, O. F., Martinez, L. J., Far, E., & Albarracin, V. H. (2019). Polyextremophilic Bacteria from High Altitude Andean Lakes : Arsenic Resistance Profiles and Biofilm Production. 2019.

95. Zenoff, V. F., Heredia, J., Ferrero, M., Siñeriz, F., & Farías, M. E. (2006). Diverse UV-B resistance of culturable bacterial community from high-altitude wetland water. Current Microbiology, 52(5), 359–362. https://doi.org/10.1007/s00284-005-0241-5

96. Zenoff, V. F., Siñeriz, F., & Farías, M. E. (2006). Diverse responses to UV-B radiation and repair mechanisms of bacteria isolated from high-altitude aquatic environments. Applied and Environmental Microbiology, 72(12), 7857–7863. https://doi.org/10.1128/AEM.01333-06

97. Zhu, D. X., Garner, A. L., Galburt, E. A., & Stallings, C. L. (2019). CarD contributes to diverse gene expression outcomes throughout the genome of Mycobacterium tuberculosis. Proceedings of the National Academy of Sciences of the United States of America, 116(27), 13573–13581. https://doi.org/10.1073/pnas.1900176116

