## Supplementary file 2 for "Proteomic Signatures of Microbial Adaptation to the Highest UV-Irradiation on Earth: Lessons from a Soil Actinobacterium"

**Supplemental figures**

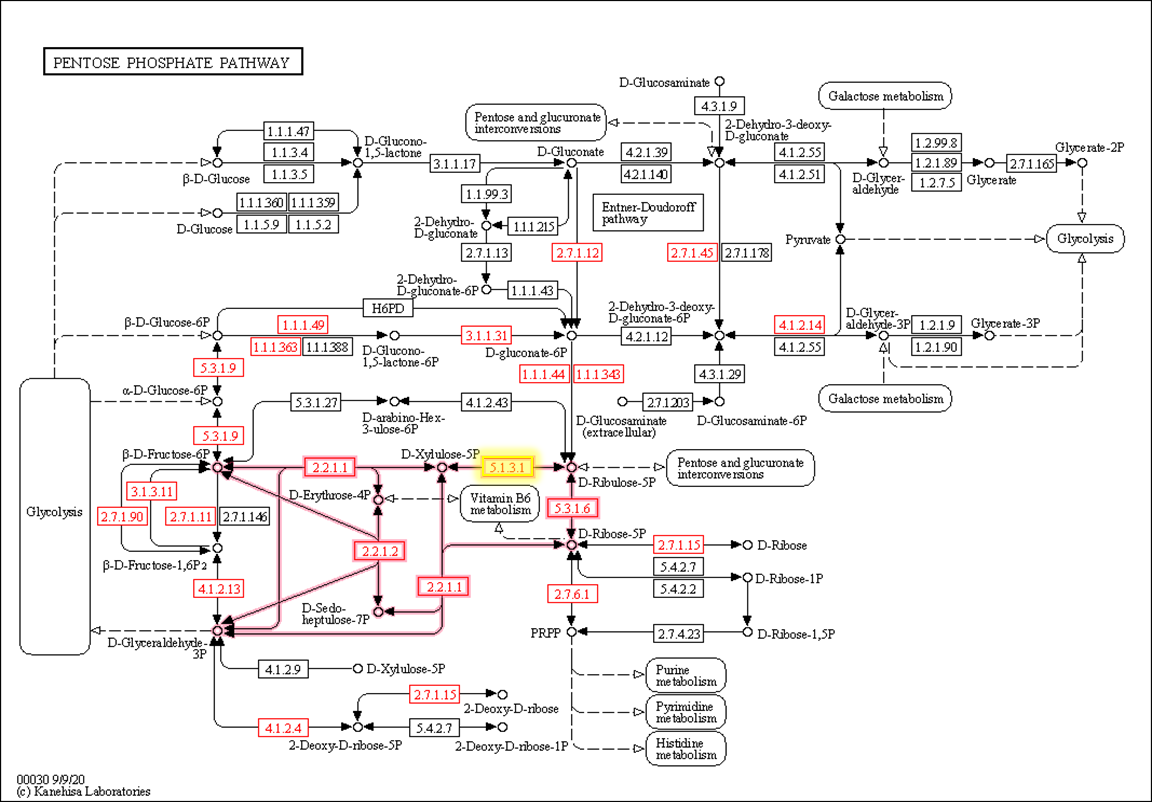

**Figure S1-** General view of pentose phosphate pathways from KEGG server. Red-marked items represent proteins encoded in *Nesterenkonia* sp. Act20 genome, whereas the yellow-marked item indicate the Ribulose-phosphate 3-epimerase (EC 5.1.3.1) (rpe) protein which was absent in Dt samples. Pink arrows indicate the non-oxidative phase of the pentose phosphate pathway.

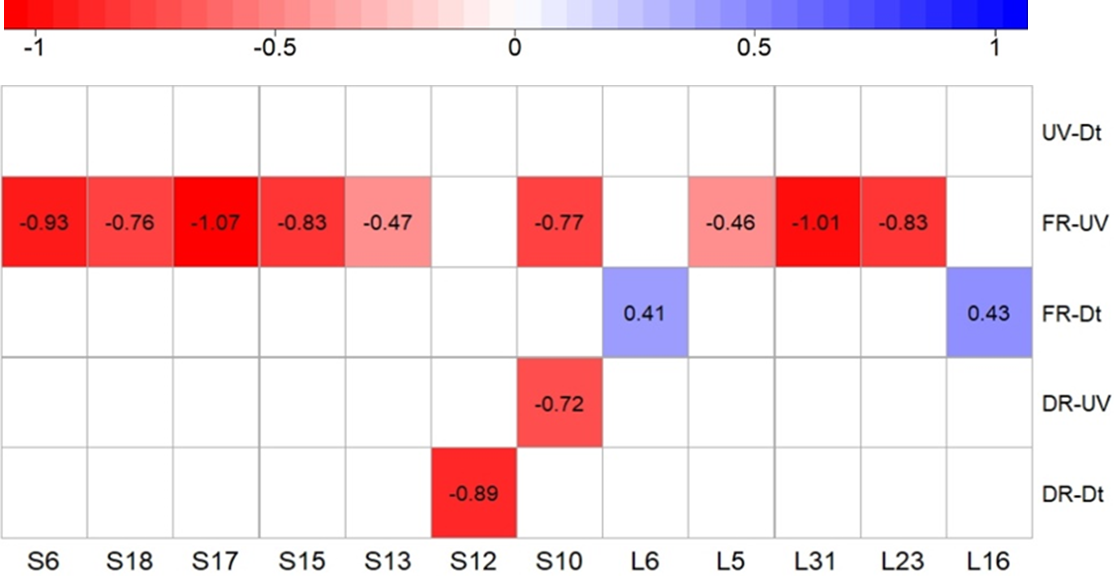

**Figura S2-** Heatmap comparing the abundance of ribosomal proteins with statistically significate values between treatments.

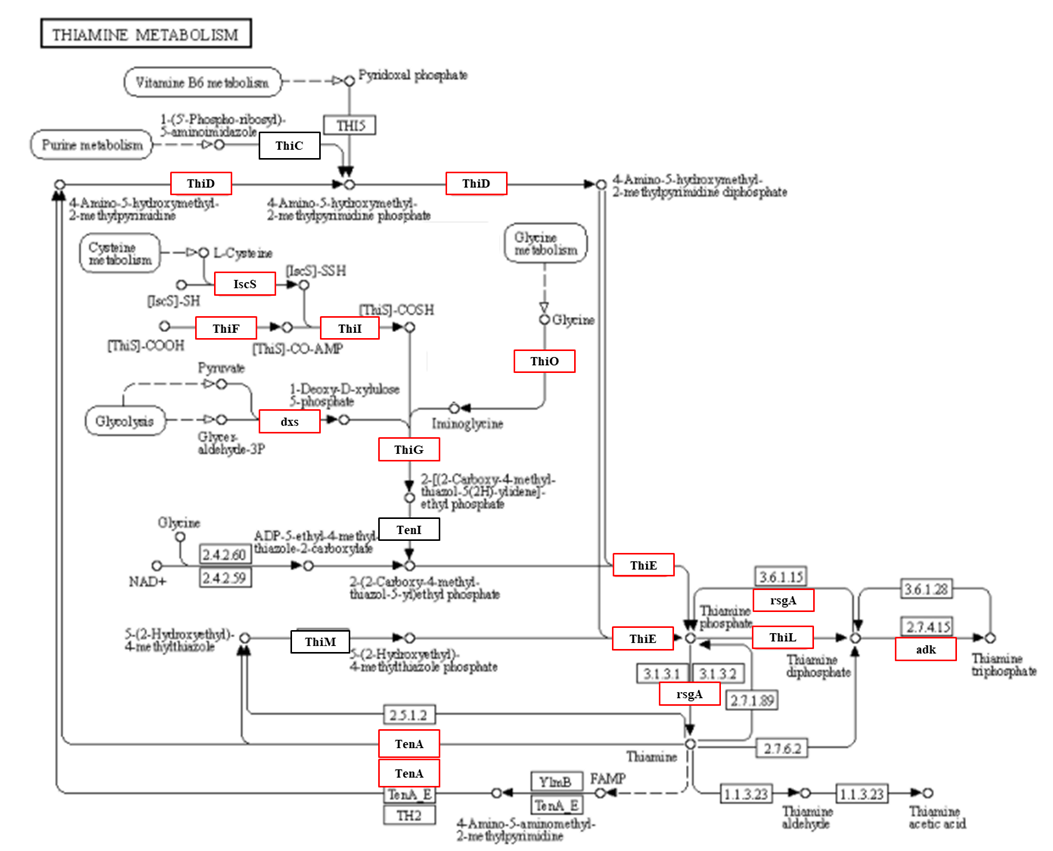

**Figura S3-** General view of thiamine biosynthesis pathways from KEGG server. Red-marked items represent proteins encoded in *Nesterenkonia* sp. Act20 genome. The EC numbers of some proteins were replaced by their canonical names in order to facilitate interpretation of readers.

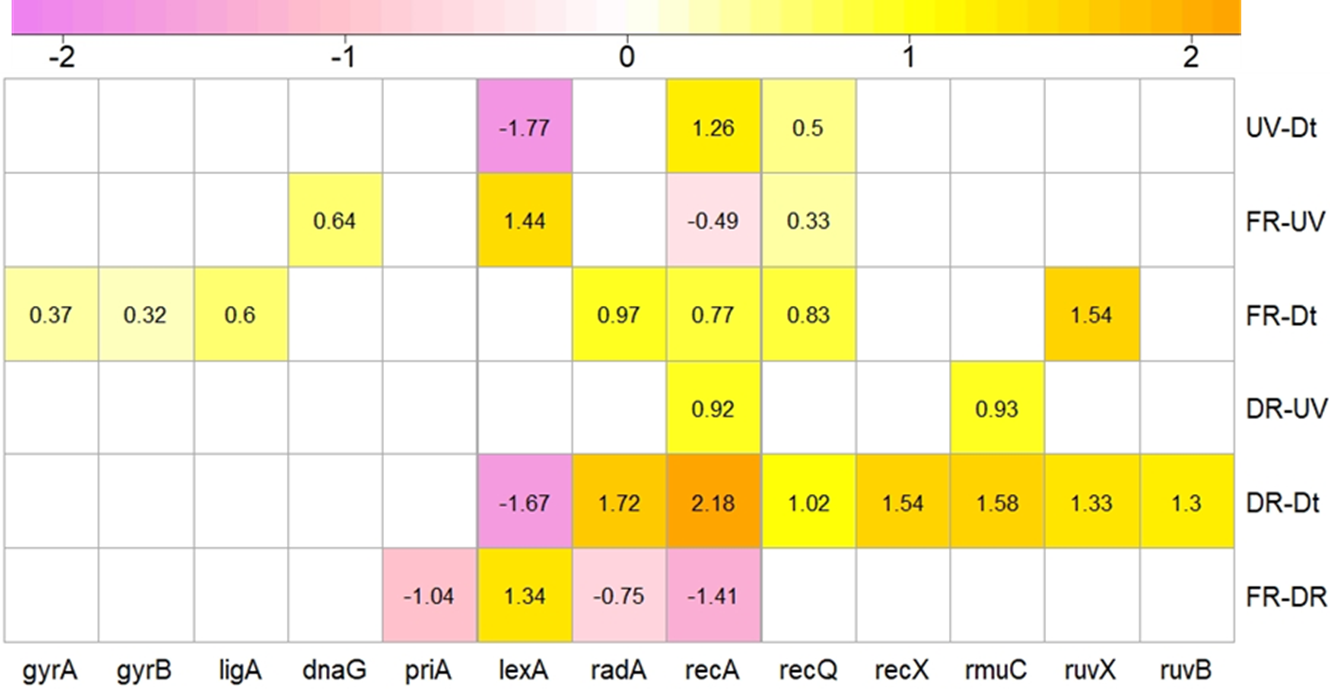

**Figure S4-** Heatmap comparing the abundance of proteins involved in homologous recombination and DNA repair between treatments.

**Note 1-** There are two definitions of "fold-change" (FC) in the literature. The standard definition, referred to as FC_ratio_ (a), is denoted as the ratio of the average values of the normalised areas of a protein for two treatments. Following this definition, a protein will be over-regulated if the average abundance value for a protein in a given treatment is greater than twice the average abundance value for the same protein in another treatment. On the other hand, FC_difference_ (b), is denoted as the difference of the mean abundance values on a logarithmic scale in base 2 of a protein for two treatments. According to this definition, a protein in a given treatment will be over-regulated relative to another treatment if the FC_difference_ value exceeds the values 1 and -1. However, FC_difference_ has advantages when the dataset is imputed as it avoids data loss by replacing invalid values with a true value.

**a)** FC*_ratio_* i = _i_ / _i_	**b)** FC*_difference_* i = _i_ - _i_

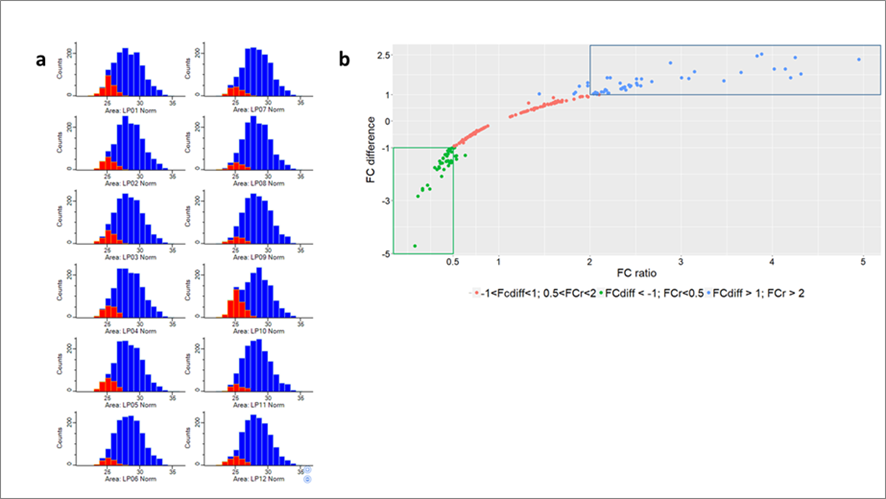

**Imputed values in proteomic dataset-** **a)** Invalid values (in red) imputed by the minimum values detected from the normal distribution of the whole dataset of each treatment for each biological replicate. **b)** Relationship between FC_ratio_ and FC_difference_ in the proteomic dataset (statistics file). It is observed that values of FC_ratio_ greater or less than twice (0.5>FC_ratio_<2) correspond to values of FC_difference_ greater than and less than 1 and -1, respectively.
